# Synaptic plasticity induced by differential manipulation of tonic and phasic motoneurons in Drosophila

**DOI:** 10.1101/2020.04.28.066696

**Authors:** Nicole A. Aponte-Santiago, Kiel G. Ormerod, Yulia Akbergenova, J. Troy Littleton

## Abstract

Structural and functional plasticity induced by neuronal competition is a common feature of developing nervous systems. However, the rules governing how postsynaptic cells differentiate between presynaptic inputs are unclear. In this study we characterized synaptic interactions following manipulations of Ib tonic or Is phasic glutamatergic motoneurons that co-innervate postsynaptic muscles at Drosophila neuromuscular junctions (NMJs). After identifying drivers for each neuronal subtype, we performed ablation or genetic manipulations to alter neuronal activity and examined the effects on synaptic innervation and function. Ablation of either Ib or Is resulted in decreased muscle response, with some functional compensation occurring in the tonic Ib input when Is was missing. In contrast, the phasic Is terminal failed to show functional or structural changes following loss of the co-innervating Ib input. Decreasing the activity of the Ib or Is neuron with tetanus toxin light chain resulted in structural changes in muscle innervation. Decreased Ib activity resulted in reduced active zone (AZ) number and decreased postsynaptic subsynaptic reticulum (SSR) volume, with the emergence of filopodial-like protrusions from synaptic boutons of the Ib input. Decreased Is activity did not induce structural changes at its own synapses, but the co-innervating Ib motoneuron increased the number of synaptic boutons and AZs it formed. These findings indicate tonic and phasic neurons respond independently to changes in activity, with either functional or structural alterations in the tonic motoneuron occurring following ablation or reduced activity of the co-innervating phasic input, respectively.

**Significance Statement:** Both invertebrate and vertebrate nervous systems display synaptic plasticity in response to behavioral experiences, indicating underlying mechanisms emerged early in evolution. How specific neuronal classes innervating the same postsynaptic target display distinct types of plasticity is unclear. Here, we examined if Drosophila tonic Ib and phasic Is motoneurons display competitive or cooperative interactions during innervation of the same muscle, or compensatory changes when the output of one motoneuron is altered. We established a system to differentially manipulate the motoneurons and examined the effects of cell-type specific changes to one of the inputs. Our findings indicate Ib and Is motoneurons respond differently to activity mismatch or loss of the co-innervating input, with the tonic subclass responding robustly compared to phasic motoneurons.

## Introduction

Functional and structural changes in neuronal circuits occur during development and in response to environmental stimuli, learning, and injury (Katz and Shatz, 1996; Destexhe and Marder, 2004; Foeller and Feldman, 2004; Lamprecht and LeDoux, 2004; Holtmaat and Svoboda, 2009). Disruptions of these plasticity pathways contribute to neurodevelopmental diseases and impair rewiring after brain injury, highlighting the importance of the underlying mechanisms (Luo and O’Leary, 2005; Melom and Littleton, 2011; Doll and Broadie, 2014; Nahmani and Turrigiano, 2014). In contrast to mammals, invertebrate nervous systems like that of *Drosophila melanogaster* are more stereotypical in their organization. Neuroblasts divide and differentiate in a specific order to generate fixed cellular lineages with genetically hard-wired synaptic targets (Hartenstein and Campos-Ortega, 1984; Thomas et al., 1984; Johansen et al., 1989a; Bossing et al., 1996; Landgraf et al., 1997; Schmid et al., 1999; Hoang and Chiba, 2001; Yu et al., 2010; Harris et al., 2015; Lee, 2017; Shepherd et al., 2019). Although Drosophila display stereotypical neuronal connectivity, plasticity can occur throughout development and into adulthood. Structural plasticity is most prominent during metamorphosis, when larval neurons reorganize their processes and synaptic partners to form functional adult circuits (Technau and Heisenberg, 1982; Truman, 1990; Schubiger et al., 1998; Lee and Luo, 1999; Marin et al., 2005; Williams and Truman, 2005; Alyagor et al., 2018; Mayseless et al., 2018). Alterations in connectivity also occur in response to changes in environmental stimuli or following acute or chronic manipulations of neuronal activity (Cash et al., 1992; Chang and Keshishian, 1996; Davis et al., 1998; Lnenicka et al., 2003; Sigrist et al., 2003; Berdnik et al., 2006; Hourcade et al., 2010; Matz et al., 2010; Golovin et al., 2019).

Although plasticity occurs broadly across neuronal circuits, the motor system has played an important role in defining mechanisms for activity-dependent structural changes in connectivity. Locomotion is an essential behavior in many animals and requires coordinated output from central pattern generators to orchestrate motoneuron firing patterns that activate specific muscles (Marder and Calabrese, 1996; Marder and Rehm, 2005). In vertebrates, individual muscle fibers receive transient innervation from many cholinergic motoneurons during early development (Sanes and Lichtman, 1999). As many as 10 distinct motor axons can innervate a single muscle fiber before an activity-dependent competition results in retention of only a single axon (Tapia et al., 2012). This axonal competition allows a large pool of identical motoneurons to transition from dispersed weak outputs to the muscle field to strong innervation of a smaller subset of muscles (Colman et al., 1997; Walsh and Lichtman, 2003; Turney and Lichtman, 2012).

Unlike vertebrate neuromuscular junctions (NMJs), early promiscuity in synaptic partner choice and subsequent synapse elimination does not occur in Drosophila. Instead, the larval motor system is comprised of ∼36 motoneurons from 4 subclasses that are genetically programmed by specific transcription factors and guidance molecules to form stereotypical connections to the 30 muscles in each abdominal hemisegment (Hoang and Chiba, 2001; Clark et al., 2018). Although synaptic partner choice is hardwired, activity-dependent plasticity and homeostatic mechanisms have been characterized, making Drosophila an ideal system to study synaptic interactions between motor neurons (Davis et al., 1998; Sigrist et al., 2003; Guan et al., 2005; Yoshihara et al., 2005; Frank et al., 2006; Berke et al., 2013; Davis, 2013; Cho et al., 2015; Davis and Müller, 2015; Gaviño et al., 2015; Harris and Littleton, 2015; Böhme et al., 2019; Goel et al., 2019). Although individual muscles normally restrict innervation to a single neuron from each subclass, it is unclear if motoneurons interact during innervation of the same muscle target or respond when the output of one motoneuron is altered. Therefore, we established a system to differentially manipulate the two primary glutamatergic inputs and characterized the subsequent effects on synaptic morphology and function. We found that only the tonic Ib motoneuron is capable of partially compensating following ablation or silencing of the phasic Is input.

## Materials and Methods

### Drosophila stocks

*Drosophila melanogaster* were cultured on standard medium at 25°C. Genotypes used in this study include: *w*^*1118*^ (BDSC 3605); 13X LexAop2-mCD8-GFP (Bloomington Drosophila Stock Center (BDSC) 32204); UAS-DsRed (BDSC 6282); UAS-CD8-GFP (BDSC 32185); UAS-Reaper (BDSC 5823); UAS-TeTXLC (BDSC 28837); UAS-NaChBac (BDSC 9469); GMR27E09-GAL4 (BDSC 49227); MN1-Ib-GAL4 (BDSC 40701); 6-58 Is-GAL4 (Ellie Heckscher, U. Chicago). Animals of either sex were used depending upon genetic scheme.

### Transgenic constructs

To create the MN1-Ib LexA driver line, a 2736 base pair fragment from the intronic region of the *Dpr4* locus that was used to generate the GMR94G06 MN1-Ib-GAL4 driver line (BDSC stock #40701) was PCR amplified and cloned into pCR8/GW/TOPO® (Invitrogen). This was followed by an LR cloning step into the pBPLexA::p65Uw plasmid (Addgene 26231). The resulting construct was sent for injection into an attP40 donor site strain by BestGene (Chino Hills, CA, USA).

### Immunocytochemistry

3^rd^ instar wandering larvae were reared at 25°C and dissected in hemolymph-like HL3.1 solution (in mM: 70 NaCl, 5 KCl, 1.5 CaCl_2_, 4 MgCl_2_, 10 NaHCO3, 5 trehalose, 115 sucrose, 5 HEPES, pH 7.2). Larvae were fixed for ten minutes in HL3.1 buffer with 4% formaldehyde and washed three times for ten minutes with PBT (PBS containing 0.1% Triton X-100), followed by a thirty-minute incubation in block solution (5% NGS in PBT). Fresh block solution and primary antibodies were added. Samples were incubated overnight at 4°C and washed with two short washes and three extended 20 minutes washed in PBT. Secondary antibodies were added to block solution and incubated at room temperature for two hours or at 4°C overnight. Finally, larvae were rewashed and mounted in 80% Glycerol. Antibodies used for this study include: mouse anti-BRP, 1:500 (NC82, Developmental Studies Hybridoma Bank (DSHB) NC82, Iowa City, IA)); mouse anti-DLG, 1:1000 (DSHB stock #4F3); chicken anti-DLG, 1:500; DyLight 649 conjugated anti-HRP, 1:500 (#123-605-021; Jackson Immuno Research, West Grove, PA, USA); rabbit anti-GFP Alexa Fluor 488, 1:500 ((#G10362, ThermoFisher Scientific, Waltham, MA, USA); goat anti-mouse Alexa Fluor 546, 1:500 (A-11030; ThermoFisher); Phalloidin-conjugated Alexa Fluor 555 or 657, 1:500 (ThermoFisher). Immunoreactive proteins were imaged on a Zeiss Pascal confocal microscope (Carl Zeiss Microscopy, Jena, GERMANY) using either a 40X 1.3 NA, 63X 1.3 NA or a 100X 1.3 NA oil-immersion objective (Carl Zeiss Microscopy). Images were processed using Zen (Zeiss) software.

### Motoneuron GAL4 driver screen

The FlyLight Project image database of larval brain and VNC GFP expression provided by Gerry Rubin (Janelia, HHMI) was searched for GAL4 lines displaying restricted expression in small subsets of segmentally repeated neurons with GFP-labeled axons projecting from the VNC. Candidate lines meeting these criteria were obtained and crossed to UAS-CD8::GFP (BDSC 32185) for immunostaining with DyLight 649 conjugated anti-HRP (Jackson ImmunoResearch), Alexa Fluor 555 Phalloidin (ThermoFisher) and rabbit anti-GFP Alexa Fluor 488 (ThermoFisher). Confocal imaging was performed to classify labeled neurons based on their synaptic connectivity within the abdominal musculature.

### Quantification of confocal images

Imaris 9.2 software (Oxford Instruments, Zurich, Switzerland) was used to identify BRP puncta to quantify AZ number, and HRP labeling was used to quantify synaptic bouton number from 3D image stacks through the NMJ. For DLG/HRP measurements, the 3D mask feature was used and the software determined bouton volume within HRP staining and muscle SSR volume from DLG staining. Quantification was conducted at muscle M1 in abdominal segment A3. The n value refers to the number of NMJs analyzed, with no more than two NMJs analyzed per larvae. Animals used in each analysis were derived from at least three independent experimental crosses. All analysis was performed blind to genotype.

### Live imaging of Ib and Is innervation and synaptic growth

Live imaging was done under desflurane anesthesia at muscles M1 and M4 at abdominal segments A2-A4 as previously described (Akbergenova et al., 2018). Selected larvae were covered with halocarbon oil and a cover glass and imaged. After imaging, larvae were placed in numbered chambers with food in a 25°C incubator. Larvae were imaged at the beginning of the 1^st^ instar larval stage and during the subsequent 24 hr interval with the same data acquisition settings. Confocal images were obtained on a Zeiss Axio Imager 2 equipped with a spinning-disk confocal head (CSU-X1; Yokagawa, Tokyo, JAPAN) and ImagEM X2 EM-CCD camera (Hamamatsu, Hamamatsu City, Japan). A pan-APOCHROMAT 63X objective with 1.40 NA from Zeiss (Carl Zeiss Microscopy) was used for imaging.

### Electrophysiology

Wandering 3^rd^ instar larvae were dissected in modified HL3.1 saline (in mM: 70 NaCl, 5 KCl, 0.3 CaCl_2_, 4 MgCl_2_, 10 NaHCO_3_, 5 Trehalose, 115 Sucrose, 5 HEPES, pH 7.18). For electrophysiology and force recordings, larvae were pinned medial side up at the anterior and posterior ends, an incision was made along the midline, and the visceral organs were removed. All nerves emerging from the CNS were severed at the ventral nerve cord, and the CNS and ventral nerve cord was removed. EJPs were elicited by stimulating severed abdominal nerves. A Master 8 A.M.P.I. stimulator (A-M Systems, Sequim, WA) was used for stimulation via a suction electrode. EJPs were recorded using sharp glass microelectrodes containing a 2:1 mixture of 3M potassium chloride:3M potassium acetate with electrode resistances of 40-80 megaohms. An Axoclamp 2B amplifier (Molecular Devices, San Jose, CA) was used for signal detection and digitized via Axon Instruments Digidata 1550 (Molecular Devices). Signals were acquired at 10 kHz using Clampex and analyzed with Clampfit, MiniAnalysis, and Microsoft Excel. All analysis was performed blind to genotype.

### Muscle force contraction measurements

Force recordings were obtained using an Aurora Scientific 403A force transducer system (Aurora Scientific, Aurora, ON, CA) with a force transducer headstage, amplifier and digitizer. Larvae were dissected ventral side up in HL3.1 saline containing 1.5 mM Ca^2+^. Nerve-evoked contractions were generated using stimulation bursts from a Master 8 stimulator (A.M.P.I.). The duration of single impulses was 5 ms and interburst duration was kept constant at 15 s. Burst frequency were altered during each individual experiment. Digitized data was acquired using Dynamic Muscle Acquisition Software (DMCv5.5, Aurora Scientific) and imported and processed in Matlab using custom code. All analysis was performed blind to genotype.

### Statistical analysis

Prism software (v. 8.1.1, GraphPad Software, La Jolla, CA, USA) and FIJI /ImageJ was used for statistical analysis. Statistical significance in two-way comparisons was determined by a Student’s t-test, while one-way ANOVA parametric analysis was used when comparing more than two datasets. Statistical comparisons are with control unless noted. Appropriate sample size was determined using a normality test. Data is presented as mean ± SEM; * = p<0.05, ** = p<0.01, *** = p<0.001, n.s. = not significant.

## Results

### Screen for Ib and Is motoneuron-specific GAL4 drivers

Four subclasses of motoneurons innervate the abdominal musculature in Drosophila, with each class defined by their synaptic partner choice, neurotransmitter or neuromodulator content, and biophysical and synaptic properties (Jan and Jan, 1976; Johansen et al., 1989a; Atwood et al., 1993; Lnenicka and Keshishian, 2000; Hoang and Chiba, 2001). Approximately 30 type Ib glutamatergic motoneurons are found per hemisegment and function as the primary driver of contraction for individual muscles. A single Ib motoneuron individually innervates a muscle fiber, allowing for fine-tuning of specific locomotor programs. The Ib neuronal subclass has big synaptic boutons with tonic release properties, containing active zones (AZs) with low release probability (*P*_*r*_) that facilitate during high frequency stimulation (Lnenicka and Keshishian, 2000; Peled and Isacoff, 2011; Melom et al., 2013; Newman et al., 2017; Akbergenova et al., 2018). Three type Is glutamatergic motoneurons per hemisegment provide input to the ventral, lateral or dorsal muscle groups, respectively. Each Is neuron innervates multiple fibers to coordinate contraction of functionally related muscles. In contrast to Ib, Is motoneurons have smaller synaptic boutons and fewer AZs with phasic release properties, including higher *P*_*r*_ release sites that undergo depression during repetitive stimulation (Figure 1A). The remaining type II and III subclasses are neuromodulatory in nature (Gorczyca et al., 1993; Stocker et al., 2018).

**Figure 1.**
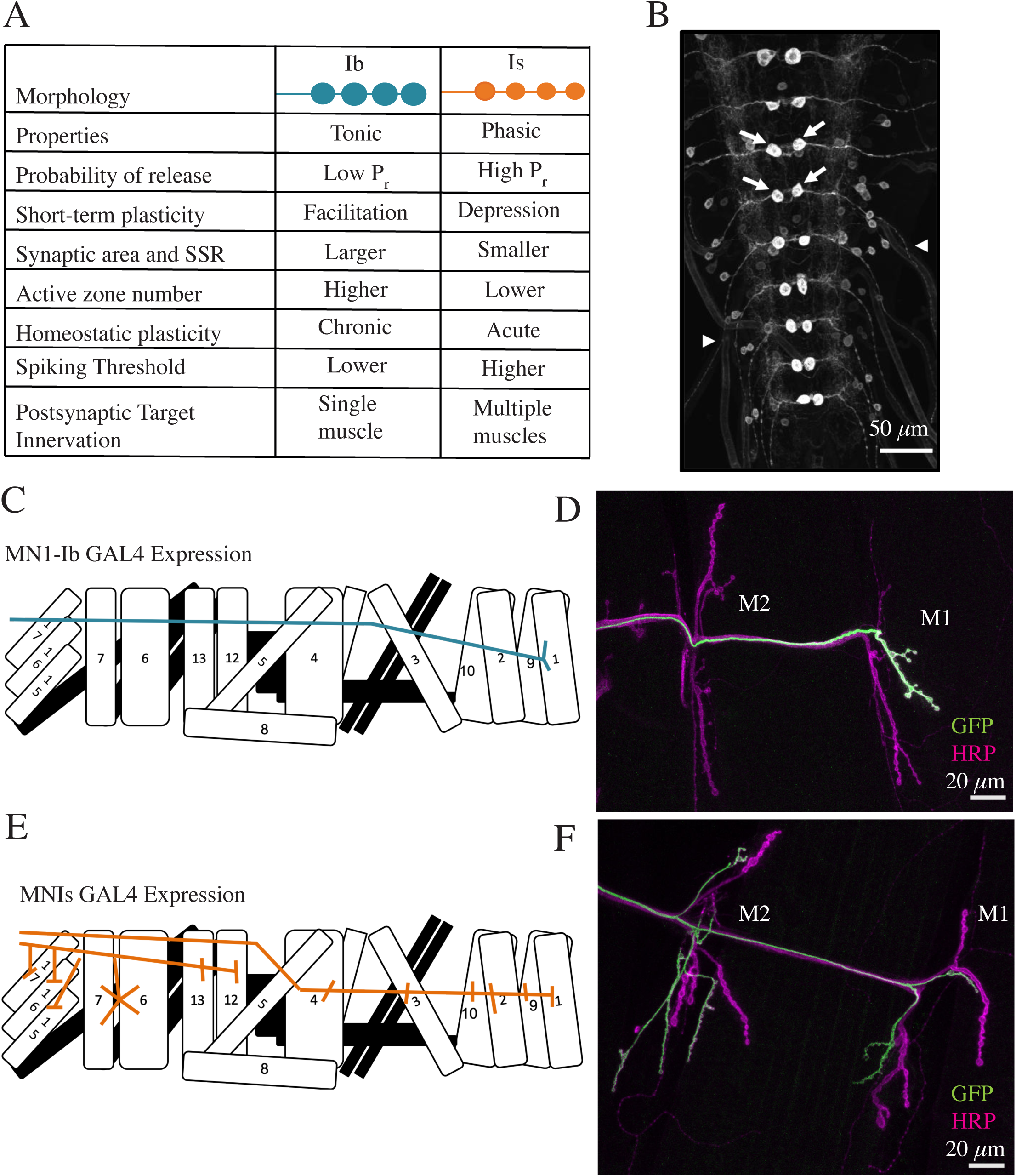
Identification of tonic Ib and phasic Is motoneuron GAL4 drivers. ***A***, Comparison of synaptic and biophysical properties of Ib and Is motoneurons in Drosophila larvae. ***B***, Confocal image of UAS-CD8-GFP driven by MN1-Ib GAL4 (GMR94G06) in the 3^rd^ instar larval VNC from the FlyLight Project GAL4 collection. Arrows denote the paired MN1-Ib cell bodies in each abdominal segment, arrowheads denote GFP expression in axons exiting the VNC. Scale bar = 50 *µ*m. ***C***, Diagram of MN1-Ib innervation in a larval abdominal hemisegment. **D**, Immunostaining for anti-GFP (green) to label MN1-Ib and HRP (magenta) to label all axons in a MN1-Ib GAL4; UAS-CD8-GFP 3^rd^ instar larva. Muscles M1 and M2 are indicated. Scale bar = 20 *µ*m. ***E***, Diagram of MNISN-Is and MNSNb/d-Is innervation in a larval abdominal hemisegment. ***F***, Immunostaining for anti-GFP (green) to label MNIs and HRP (magenta) to label all axons in a MNIs GAL4 (6-58); UAS-CD8-GFP 3^rd^ instar larva. Muscles M1 and M2 are indicated. Scale bar = 20 *µ*m.

To preferentially manipulate Ib and Is motoneurons, GAL4 lines with subclass-specific expression were identified from the FlyLight Project (Jenett et al., 2012; Manning et al., 2012) and strains provided by Ellie Heckscher (Univ. of Chicago). The FlyLight collection consists of >5000 transgenic Drosophila lines with ∼3 kb of regulatory genomic DNA from candidate neuronal genes driving GAL4 expression. Images of membrane-tethered UAS-CD8-GFP driven by each GAL4 line in 3^rd^ instar brain lobes and ventral nerve cord (VNC) were provided by Gerry Rubin (Janelia, HHMI). Candidate lines for further analysis were chosen on the basis of two criteria: (1) restricted expression of GFP in a single pair or small subset of segmentally repeated abdominal neurons in the VNC; and (2) GFP expression in axons exiting the VNC as expected for motoneurons innervating peripheral musculature (Figure 1B). Forty-two GAL4 driver lines were identified as promising candidates in the initial screen and subjected to further immunostaining to identify their synaptic targets. Six of these lines were verified as having restricted expression in a small subset of motoneurons (Table 1), including GAL4 drivers specific for Ib and Is motoneurons (Figure 1C-F). Line GMR94G06 displayed restricted expression in the tonic Ib motoneuron (MN1-Ib) that innervates muscle M1 (Figure 1B-D). GMR94G06 contains regulatory DNA from the *Dpr4* locus, which encodes a member of the cell surface Ig-containing proteins implicated in synaptic target recognition (Carrillo et al., 2015). Line GMR27F01 contained regulatory sequences from the *Fmr1* gene and showed restricted expression in two Is motoneurons and a type II neuromodulatory neuron (Table 1). Line 6-58 displayed restricted expression in the Is motoneurons MNISN-Is and MNSNb/d-Is that innervate the ventral and dorsal muscles, respectively (Figure 1E-F). To determine the gene regulatory region responsible for Is motoneuron expression in 6-58, which contained an unknown insertion site, plasmid rescue and reverse PCR was performed. The insertion site mapped to the 5’ UTR of the *Dip-α* gene. Like DPR4, DIP-α is a member of the Ig domain family of synaptic target recognition proteins and was previously found to be expressed in Is motoneurons (Ashley et al., 2019). The MN1-Ib driver GMR94G06 and an additional Is GAL4 driver line (GMR27E09, containing regulatory sequences from *Fmr1*) were independently identified in a recent study (Pérez-Moreno and O’Kane, 2019). We refer to the restricted GAL4 driver lines as MN1-Ib GAL4 and MNIs GAL4 in the remaining text. Together, they provide a toolkit to genetically manipulate the tonic Ib and phasic Is motoneuron subclasses that co-innervate muscle M1.

**Table 1.**
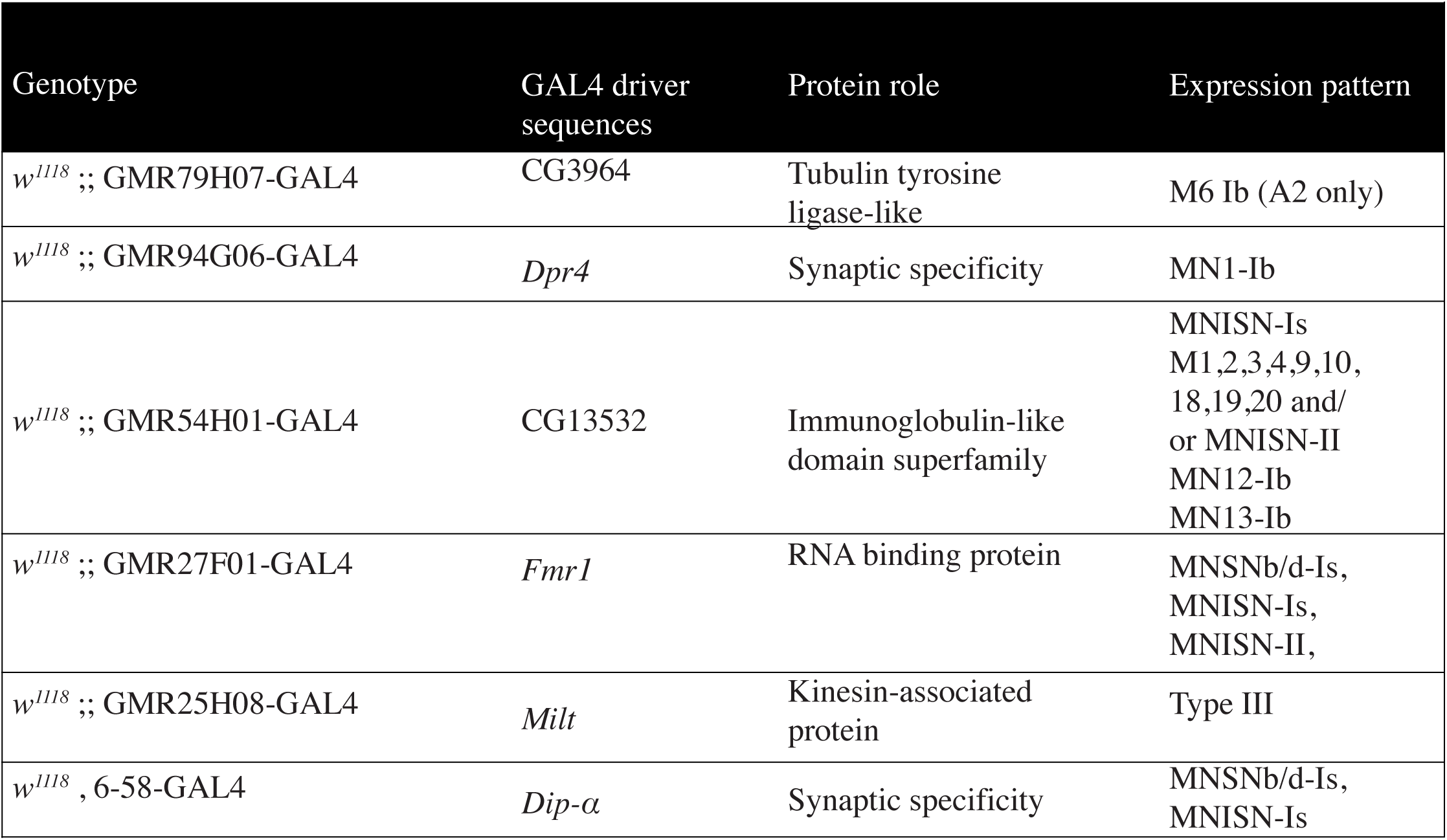
The genotype, GAL4 driver enhancer sequence or site of insertion, predicted protein role and expression pattern revealed by crossing to UAS-CD8-GFP for 3^rd^ instar larvae is shown for the six lines identified with motoneuron-restricted expression. GMR94G06 and 6-58 are referred to as MN1-Ib GAL4 and MNIs GAL4, respectively, throughout the manuscript.

### Establishment of MN1-Ib and MN-Is synaptic connections during development

MN1-Ib and MNIs drivers were used to fluorescently label the two neuronal subpopulations and establish the timing for M1 innervation, the most peripheral target of the dorsal abdominal musculature. MN1-Ib (formerly referred to as aCC) and MNIs (formerly referred to as RP2) were previously identified as pioneer neurons for the ISN nerve branch, being the first to exit the VNC towards the dorsal muscle field (Jacobs and Goodman, 1989; Johansen et al., 1989b; Lin et al., 1995; Sánchez-Soriano and Prokop, 2005). To co-label MN1-Ib and MNIs in the same animal, a MN1-Ib LexA driver line was generated by subcloning the 2736 bp genomic *Dpr4* fragment from GMR94G06 into pBPLexA::p65Uw. This construct was used to generate transgenic animals containing MN1-Ib LexA, allowing independent LexA and GAL4 transgene expression in MN1-Ib and MNIs motoneurons. Serial intravital imaging through the cuticle of briefly anesthetized animals co-expressing MN1-Ib Lex; LexAop2-CD8-GFP and MNIs GAL4; UAS-DsRed was performed as previously described (Akbergenova et al., 2018). By the beginning of the 1^st^ instar larval stage, all MN1-Ib motoneurons had correctly targeted M1 during late embryonic development, elaborating a growth-cone like projection over the muscle surface (Figure 2A, B). In contrast, MNIs had a more variable time-course of innervation, with the Is growth cone trailing behind MN1-Ib, often without targeting M1 in early 1^st^ instars. By the end of the 1^st^ instar larval stage, only 28% of MNIs motoneurons had innervated the muscle. MNIs innervation of M1 continued over the rest of larval development, with 72% of Is motoneurons innervating M1 by the 3^rd^ instar stage (abdominal segments 2-4, n=7 larvae, Figure 2C). The remaining M1 muscles lacked Is innervation. These data indicate the tonic Ib motoneuron innervates M1 prior to the arrival of Is, with the phasic motoneuron forming synaptic contacts with the muscle later in development or failing to innervate the target completely.

**Figure 2.**
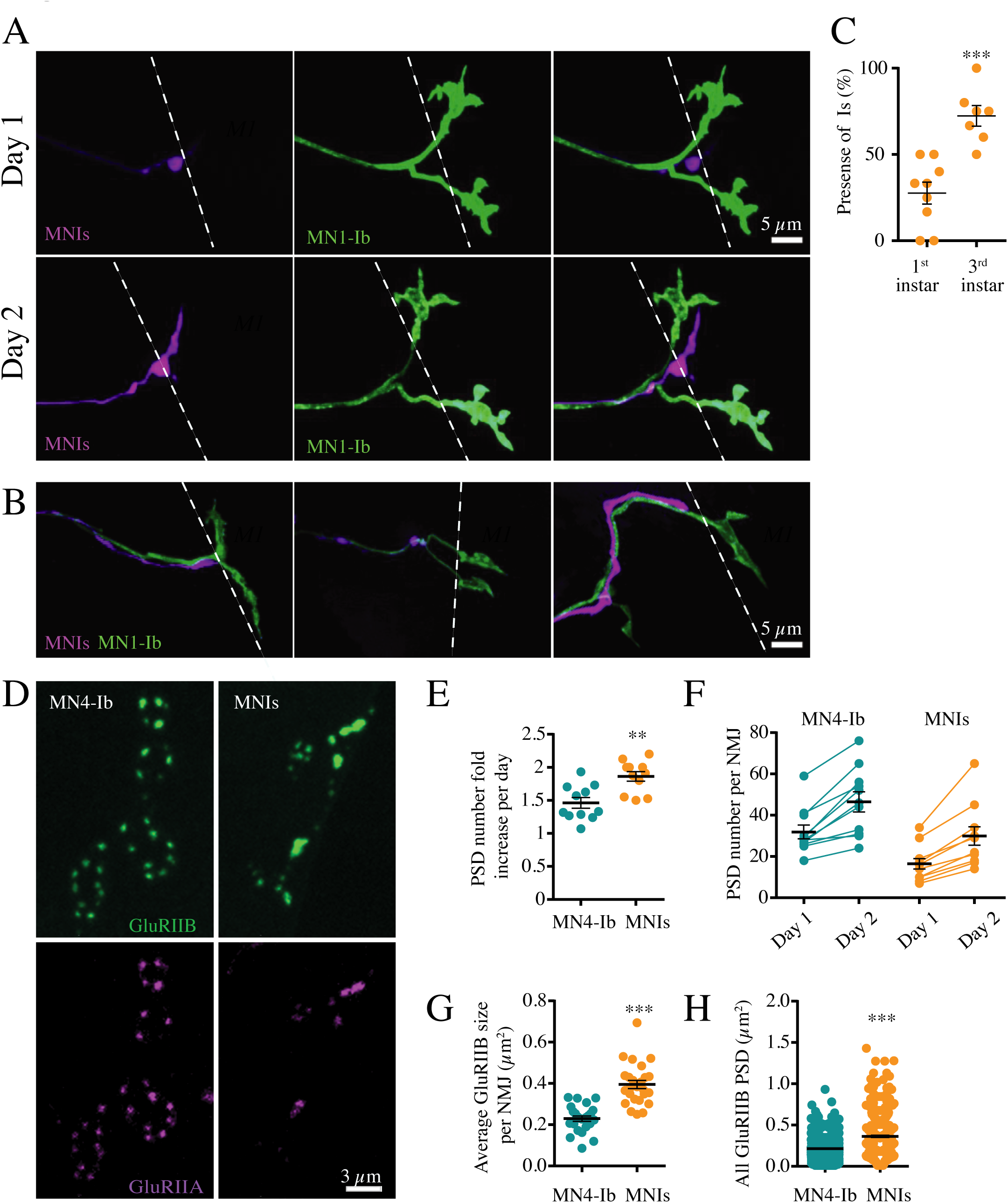
Quantification of MN1-Ib and MNIs target innervation and synapse formation with serial intravital imaging across development. ***A***, Sequential confocal images of muscle M1 innervation by MN1-Ib (green) and MNIs (magenta) at day 1 (top panels) and day 2 (bottom panels) of larval development in dual-labeled animals (MN1-Ib LexA>LexAop2-CD8-GFP; MNIs GAL4>UAS-DsRed). Dashed line indicates M1 muscle boundary. MNIs has delayed innervation compared to MN1-Ib. Scale bar = 5 *µ*m. ***B***, Representative confocal images of three M1 muscles on day 1 showing delayed innervation by MNIs (magenta) compared to MN1-Ib (green). MNIs axons in the left and middle panels proceeded to innervate M1 later in development, while the MNIs axon on the right failed to innervate M1. Dashed line indicates M1 muscle boundary. Scale bar = 5 *µ*m. ***C***, Quantification of Is motoneuron innervation of M1 in 1^st^ instar (27.6 ± 6.3%, n=9 larvae) versus 3^rd^ instar (72.4 ± 6.1%, n=7 larvae, p=0.0002, Student’s t-test). Each point represents the average percent of M1 innervation in segments A2-A4 from a single larva. ***D***, Confocal imaging of PSDs formed at MN4-Ib and MNIs NMJs on M4 in larvae expressing RFP-tagged GluRIIA (magenta) and GFP-tagged GluRIIB (green). Note the Is terminal has fewer synapses but larger PSDs. Scale bar = 3 *µ*m. ***E***, Increase in GluRIIB-positive PSDs over 24 hrs starting at the 1^st^ instar larval stage. The increase in PSD number is plotted as fold-increase of day 2 PSDs over the initial day 1 PSDs for MN4-Ib and MNIs. Each point represents the average increase at M4 from segments A2-A4 for a single larva. ***F***, Increase in PSD number at M4 during serial imaging of MN4-Ib and MNIs over 24 hrs beginning at the 1^st^ instar stage. Each point represents the average PSD number at M4 from segments A2-A4 for a single larva on day 1 and day 2. ***G***, Quantification of GluRIIB-positive PSD area for MN4-Ib and MNIs synapses at M4 in 1^st^ instar larvae. Each point represents the average PSD area at M4 from segments A2-A4 for a single larva. ***H***, Distribution of individual PSD sizes at M4 from segments A2-A4 for MN4-Ib and MNIs NMJs for all 1^st^ instar larvae imaged (n=9 larvae each). Statistical significance was determined using Student’s t-test. Data is shown as mean ± SEM; ** = p<0.01, *** = p<0.001.

Since phasic Is motoneurons have stronger synapses with higher *P*_*r*_ active zones than their Ib counterparts at the 3^rd^ instar stage (Kurdyak et al., 1994; Lnenicka and Keshishian, 2000; Lu et al., 2016; Newman et al., 2017; Genç and Davis, 2019; Karunanithi et al., 2020), we examined Is synaptic maturation given their shorter developmental time window compared to the pre-existing Ib input. Presynaptic bouton and AZ number at motoneuron NMJs rapidly increase throughout larval development, accompanied by expansion in the size of the postsynaptic density (PSD) and glutamate receptor fields (Zito et al., 1999; Akbergenova et al., 2018). To examine glutamate receptor field formation and maturation at developing tonic and phasic synapses, we followed muscle 4 (M4) innervation during early larval development (Figure 2D). The Ib and Is inputs arrive at distinct positions on M4, allowing unambiguous identification of neuronal subclass without having to genetically label the motoneurons as required for M1. Live imaging was performed in developing larvae expressing RFP-tagged GluRIIA and GFP-tagged GluRIIB under the control of their endogenous promotors (Rasse et al., 2005). Glutamate receptors at the NMJ are tetramers, with three essential subunits and a 4^th^ subunit of either GluRIIA or GluRIIB (Schuster et al., 1991; Petersen et al., 1997; Marrus et al., 2004; Featherstone et al., 2005; Qin et al., 2005). As observed at M1, Ib innervation preceded Is arrival at M4, with 18% of M4 fibers completely lacking Is innervation at the 3^rd^ instar stage (abdominal segments A2-A4, n=9 larvae). Although Is innervation of M4 was delayed compared to Ib, the rate of synapse formation, quantified by appearance of new GluRIIA/GluRIIB positive PSDs, was increased at Is terminals during consecutive days of imaging (MN4-Ib: 1.46-fold increase, MNIs: 1.86-fold increase, n=11, p=0.0014, Figure 2E, F). Overall, the delayed innervation by Is resulted in reduced PSD number at M4 compared to Ib (day 1: MN4-Ib, 31.9 ± 3.3 AZs; MNIs, 16.5 ± 2.6 AZs; day 2: MN4-Ib, 46.5 ± 4.9 AZs; MNIs, 29.9 ± 4.5 AZs, n=11, p=0.0014, Figure 2F). Although the Is motoneuron formed fewer synapses than Ib, the average PSD size defined by GluRIIB area in 1^st^ instar larvae was 69% larger than those of the corresponding Ib input (MN4-Ib: 0.215 ± 0.004 µm^2^, n=1006 PSDs; MNIs: 0.363 ± 0.013 µm^2^, n=399 PSDs, p=1.2×10^−8^, Figure 2G, H). Given PSD maturation is activity-dependent at NMJs (Schmid et al., 2008; Petzoldt et al., 2014; Akbergenova et al., 2018), these data suggest PSDs may develop faster at the stronger Is AZs than PSDs apposed to the weaker Ib AZs. Alternatively, the postsynaptic muscle may compartmentalize delivery of PSD material to Is versus Ib synapses such that Is sites are favored.

A central pathway regulating synaptic maturation at Drosophila NMJs is mediated through muscle secretion of Glass Bottom Boat (Gbb), a BMP ligand that acts on presynaptic BMP receptors to activate a SMAD-dependent transcriptional synaptic growth program (Aberle et al., 2002; Marqués et al., 2002; McCabe et al., 2003; Ball et al., 2010; Rodal et al., 2011; Berke et al., 2013). To determine if tonic Ib and phasic Is motoneurons were equally sensitive to Gbb signaling given their different innervation time course, we assayed synapse formation and growth of MN4-Ib and MNIs at M4 in *Gbb* mutants (Figure 3A). As previously observed, loss of Gbb reduced synaptic growth of Ib motoneurons innervating M4 compared to controls (control MN4-Ib NMJ area: 182.8 ± 21.6 µm^2^, n=11 NMJs from 5 larvae; *Gbb* MN4-Ib NMJ area: 56.2 ± 5.8 µm^2^, n=21 NMJs from 8 larvae, p=0.0001, Figure 3B). Although synaptic growth was reduced, 100% of M4 muscles displayed Ib synaptic innervation. In contrast, synaptic innervation from the phasic Is motoneuron was reduced in *Gbb* mutants, with only 48% of M4 muscles containing Is innervation compared to 82% in controls (Figure 3C). In cases where the Is motoneuron innervated M4 in *Gbb*, similar reductions in synaptic growth compared to Ib were observed (control Is NMJ area: 102.2 ± 11.3 µm^2^, n=7 NMJs from 5 larvae; *Gbb* Is NMJ area: 33.8 ± 4.0 µm^2^, n=16 NMJs from 8 larvae, p=0.0006, Figure 3B). These data indicate the Gbb pathway promotes synaptic growth in tonic Ib motoneurons but is not required for target innervation. In contrast, loss of Gbb signaling in phasic Is motoneurons decreases synaptic growth and also reduces the percent of muscles with Is innervation.

**Figure 3.**
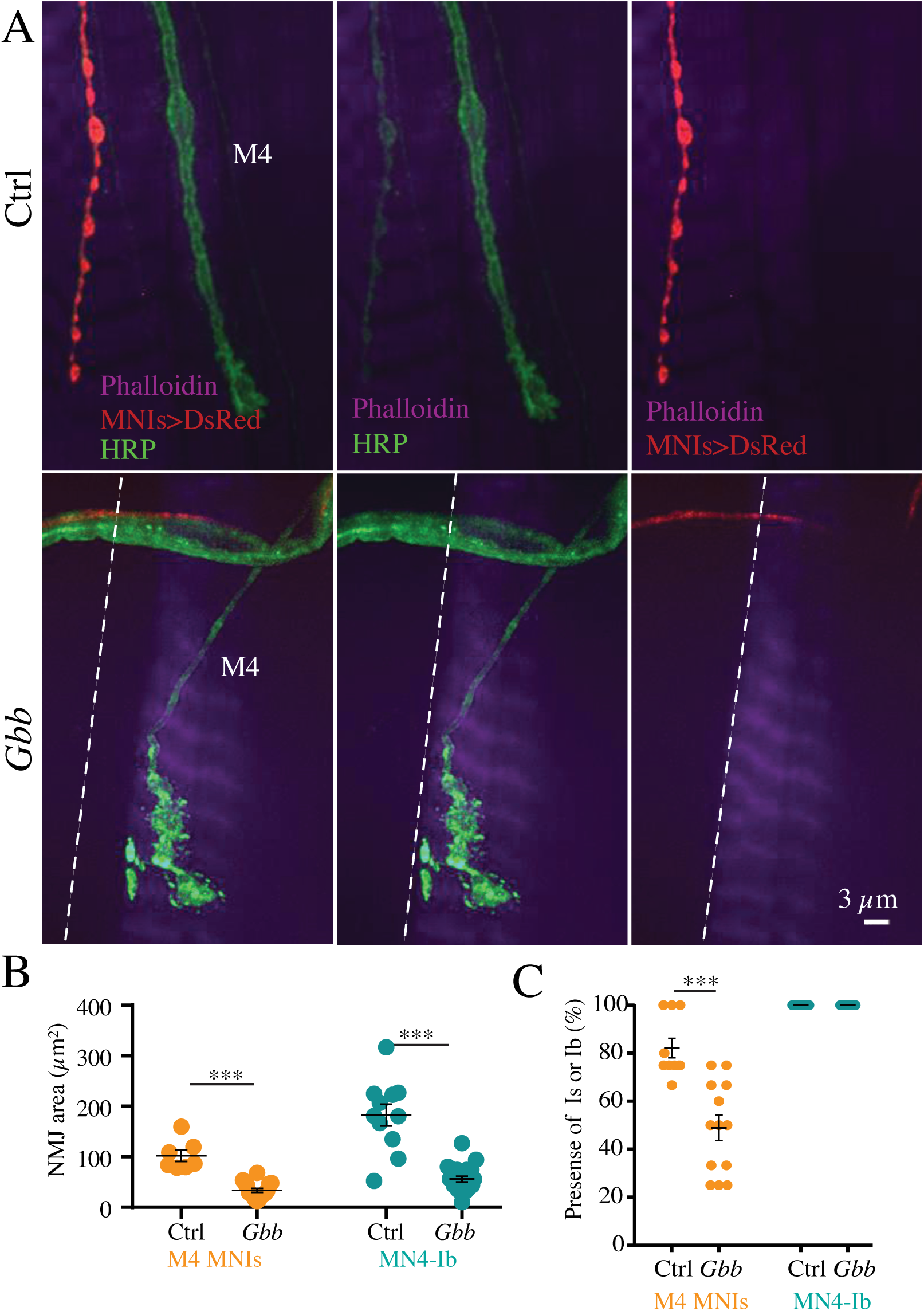
Reduction in synaptic growth and muscle innervation by Is motoneurons in *Gbb* mutants. ***A***, Confocal images of 3^rd^ instar muscle M4 innervation by MN4-Ib (green, anti-HRP staining) and MNIs (magenta, MNIs GAL4>UAS-DsRed) in controls (top panels) or *Gbb* mutants (*gbb*^*1*^*/gbb*^*2*^). Muscles were labeled with Phalloidin-conjugated Alexa Fluor 647. Dashed line indicates M4 muscle boundary. Scale bar = 3 *µ*m. ***B***, Quantification of NMJ area for MNIs and MN4-Ib at 3^rd^ instar M4 defined by anti-HRP staining for controls and *Gbb* mutants. Each point represents the average NMJ area at M4 from segments A2-A4 for a single larva. ***C***, Quantification of the percent of M4 muscles innervated by MNIs or MN4-Ib at the 3^rd^ instar stage. Each point represents the average percent of M4 innervation in segments A2-A4 from a single larva. Statistical significance was determined using Student’s t-test. Data is shown as mean ± SEM; *** = p<0.001.

### Role of MN1-Ib and MNIs in muscle excitability and contraction

To determine the relative contribution of Ib and Is motoneurons in muscle excitability, simultaneous electrophysiological recordings were performed at 3^rd^ instar larval muscles M1 and M2 in HL3.1 saline containing 0.3 mM extracellular Ca^2+^. A minimal stimulation protocol was used to isolate MN1-Ib or MNIs, as MNIs innervates both muscles compared to MN1-Ib (Figure 4A). By increasing current applied to the larval nerve through the stimulating electrode, responses following activation of one or both motor axons could be isolated in cases where dual innervation was present. The average excitatory junctional potential (EJP) amplitude recorded at M1 when both Ib and Is inputs were active was 24.2 ± 1.7 mV (n=22, Figure 4B, C). When Ib or Is were individually recruited during minimal stimulation, reduced responses of similar amplitude at M1 were observed (MN1-Ib: 11.8 ± 1.6 mV, n=22; MNIs: 12.4 ± 1.4 mV, n=22), indicating each neuron provides similar drive to M1 following single action potentials (Figure 4B, C). Given MN1-Ib has more AZs compared to MNIs at M1 (Figure 2F), these results are consistent with MNIs motoneurons having higher *P*_*r*_ per AZ as previously described (Lu et al., 2016; Newman et al., 2017; Genç and Davis, 2019).

**Figure 4.**
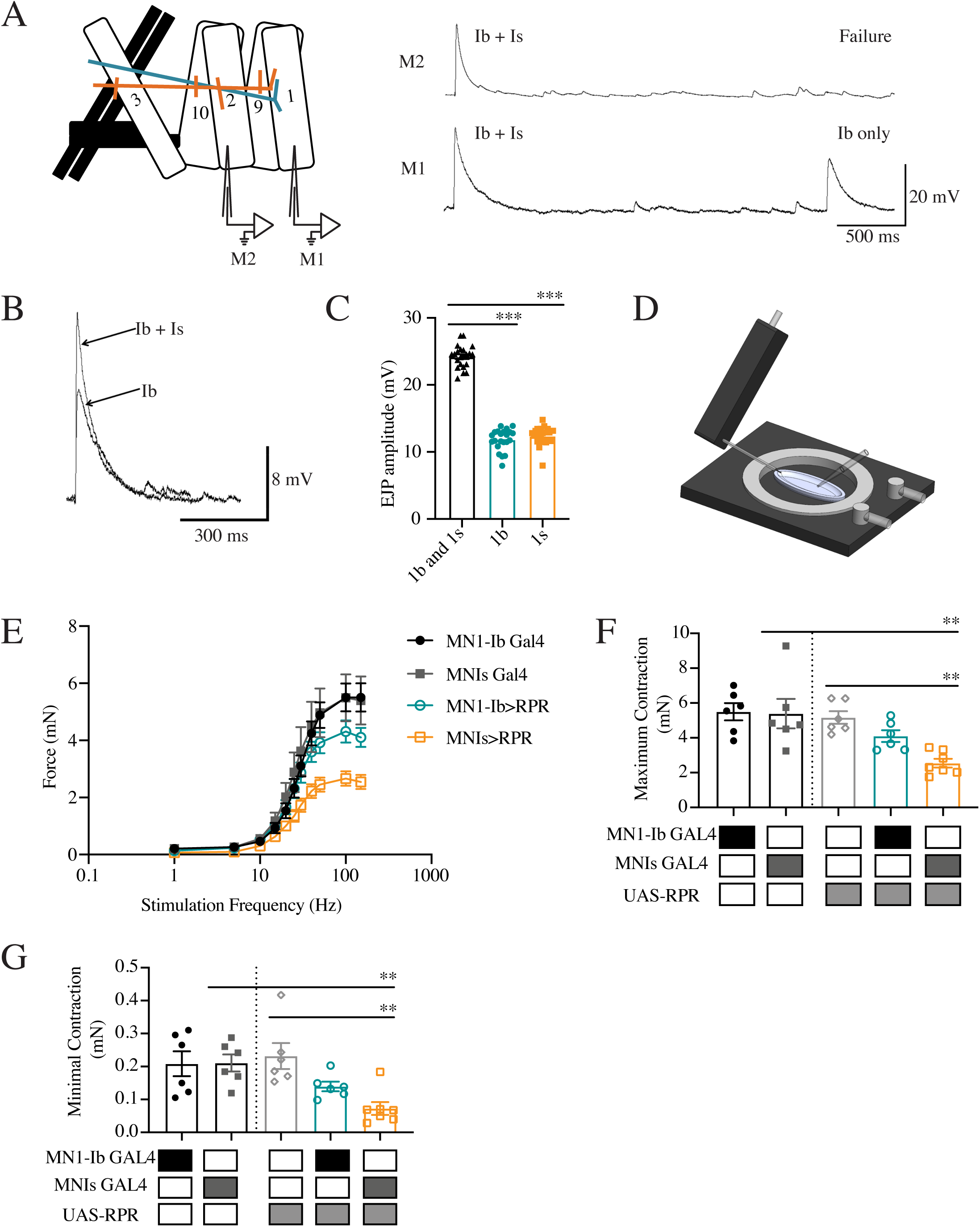
Contributions of MN1-Ib and MNIs to muscle excitability and contractile force. ***A***, Depiction of dual intracellular electrode paradigm for performing simultaneous voltage recordings from muscles M1 and M2 in control *w*^*118*^ 3^rd^ instar larvae, with MN1-Ib (teal) and MNIs (orange) labeled. Representative recordings from M1 and M2 are shown on the right. Ib + Is shows the compound EJP generated when both motoneurons are activated. Lowering stimulation intensity results in recruitment of only MN1-Ib or MNIs. Stimulation of only MNIs triggers responses in both muscles given it innervates M1 and M2. Stimulation of MN1-Ib, as shown in the Ib-only trace, results in responses only from M1. ***B***, Representative traces of simple or compound EJPs at M1 showing recruitment of MN1-Ib only or both MN1-Ib and MNIs. ***C***, Average EJP amplitude at M1 following recruitment of both motoneurons or MN1-Ib or MNIs only (n=22 larvae). ***D***, Schematic of force transducer setup used to measure larval muscle contractile force. ***E***, Force-frequency curves for 1 Hz to 150 Hz nerve stimulation in MN1-Ib and MNIs GAL4 controls and MN1-Ib GAL4>RPR and MNIs GAL4>RPR ablated 3^rd^ instar larvae. Six replicate contractions were generated at each stimulation frequency for each recording and averaged across 7 larvae per genotype. ***F***, Maximal contraction force elicited at 150 Hz is shown. Shaded boxes under each bar indicate the genotypes for each experimental group. ***G***, Minimal contraction force elicited by a single action potential for the indicated genotypes. Statistical significance was determined using Student’s t-test. Data is shown as mean ± SEM; * = p<0.01, *** = p<0.001.

To examine the contribution of MN1-Ib and MNIs for muscle contractility, nerve-evoked bodywall contraction force was recorded with a force transducer attached to the head of dissected larvae (Figure 4D). To isolate contraction force mediated predominantly by the dorsal muscle group (M1, M2, M3, M9, M10), 3^rd^ instar larvae were dissected along the ventral midline. Severed abdominal nerves were placed in a suction electrode and the nerve bundle was stimulated at increasing frequencies in HL3.1 saline containing 1.5 mM extracellular Ca^2+^ as previously described (Ormerod et al., 2016). Control MN1-Ib and MNIs GAL4 larval preparations showed increasing contraction force when a 0.1 msec simulation was ramped from 1 Hz to 150 Hz for 600 ms (Figure 4E). To determine the contribution of each motoneuron subclass for contractile force, MN1-Ib or MNIs was ablated by expressing the cell death gene *reaper* (UAS-RPR) to induce apoptosis (White et al., 1994, 1996; Goyal et al., 2000). Expression of UAS-RPR with MN1-Ib or MNIs GAL4 resulted in elimination of the corresponding motoneuron class. Ablation of MNIs removed phasic input to all dorsal and ventral muscles and resulted in a robust reduction in contractile force over the entire frequency distribution, including a 53% decrease in maximum force following 150 Hz stimulation (MNIs GAL4: 5.4 ± 0.2 mN, n=7; MNIs>UAS-RPR: 2.5 ± 0.1 mN, n=8, p=0.009, Figure 4F) and a 65% decrease in minimal contractile force following a single action potential (MNIs GAL4: 0.2 ± 0.02 mN, n=7; MNIs>RPR: 0.07 ± 0.02 mN, n=7, p=0.0009, Figure 4G). Ablation of MN1-Ib eliminated tonic input only to M1, leaving innervation of the other dorsal muscles by their respective Ib and Is neurons intact. Loss of MN1-Ib caused a less severe defect, resulting in a 25% decrease in contractile force at 150 Hz (MN1-Ib GAL4: 5.5 ± 0.2 mN, n=7; MN1-Ib>UAS-RPR: 4.1 ± 0.2 mN, n=7, p=0.022, Figure 4F) and a 33% decrease at 1 Hz (MN1-Ib GAL4: 0.21 ± 0.02 mN, n=7; MN1-Ib>UAS-RPR: 0.14 ± 0.01 mN, n=7, p=0.068, Figure 4G). These results indicate both motoneuron subclasses contribute to muscle contraction force, with the phasic Is input providing major drive for both excitability and contraction.

### Lack of Ib and Is synaptic competition during NMJ development

Given the role of tonic and phasic motoneuron inputs in driving muscle excitability, we examined whether interactions between the inputs occurred during larval development that shaped their axonal arbor expansion and AZ number when they co-innervated M1 or M4. If Ib and Is neurons competed for synaptic growth signals emanating from the muscle, or suppressed growth of the co-innervating input, competitive interactions should generate a negative correlation between Ib and Is synapse number. If the two inputs display cooperative interactions during growth, for example by co-activating the muscle to release more synaptogenic factors, one would expect a positive correlation. Similarly, if both inputs were independent and growing only in response to muscle size, a positive correlation would be expected. Ib and Is synaptic terminals were identified following anti-Discs large (DLG) immunostaining. DLG is a component of the postsynaptic muscle SSR and prominent around presynaptic Ib boutons compared to Is (Lahey et al., 1994). Synapse number was quantified for wandering 3^rd^ instar larvae by immunolabeling for the core AZ T-bar component Bruchpilot (BRP) (Wagh et al., 2006; Fouquet et al., 2009). Synaptic bouton number was determined using anti-HRP immunostaining, which provides a neuron-specific membrane label. No correlation was observed between Ib and Is AZ number per NMJ at M1 (r = -0.11, n=29, p=0.57, Pearson R, Figure 5A) or between Ib and Is inputs at M4 (Pearson R = -0.10, n=19, p=0.69, Figure 5B). Similarly, no correlation for Ib versus Is bouton number at M1 (Pearson R = 0.15, n=29, p=0.44, Figure 5C) or M4 (Pearson R = 0.13, n=19, p=0.60, Figure 5D) was found. These observations are consistent with Ib and Is motoneurons forming synapses on the muscle without obvious competitive or cooperative interactions that shape their connectivity during development.

**Figure 5.**
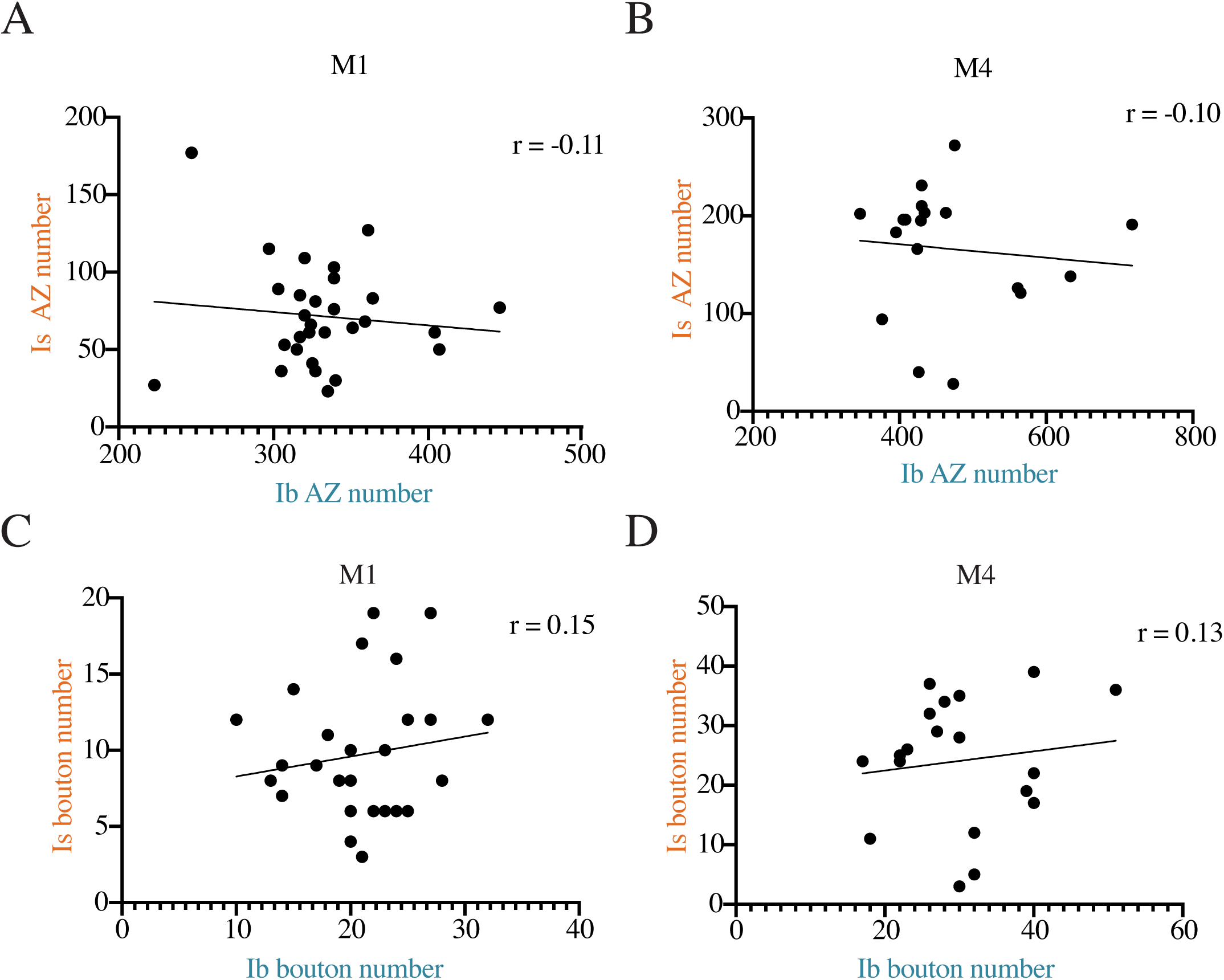
Lack of correlation between Ib and Is synaptic innervation at M1 and M4. ***A***, Correlation of MN1-Ib and MNIs AZ number at M1 quantified following immunolabeling for BRP in control *w*^*118*^ 3^rd^ instar larvae (r=-0.11, n=29, p=0.57). ***B***, Correlation of MN4-Ib and MNIs AZ number at M4 quantified following immunolabeling for BRP in *w*^*118*^ 3^rd^ instar larvae (r=-0.10, n=19, p=0.69). ***C***, Correlation of MN1-Ib and MNIs synaptic bouton number at M1 quantified following immunolabeling for HRP in *w*^*118*^ 3^rd^ instar larvae (r=0.15, n=29, p=0.44). ***D***, Correlation of MN4-Ib and MNIs synaptic bouton number at M4 quantified following immunolabeling for HRP in *w*^*118*^ 3^rd^ instar larvae (r=0.13, n=19, p=0.60). The Pearson correlation coefficient (r) is shown on the upper right for each analysis. Each data point corresponds to Ib and Is AZ or bouton number from a single larva at the indicated muscle of segment A3.

As described above, ∼30% of larval M1 muscles lack Is innervation. This natural variation provided an opportunity to examine if growth of the co-innervating MN1-Ib motoneuron was altered at mature 3^rd^ instar NMJs when MNIs innervation was absent. Lack of Is innervation did not result in any change in M1 muscle surface area (co-innervation: 46661 ± 1457 µm^2^, n=16; Ib only: 48206 ± 1170 µm^2^, n=8, p=0.66, ANOVA). Although total AZ number was reduced at M1 in the absence of Is innervation (Ib only: 285.8 ± 29.7 AZs, n=10; co-innervation: 389.7 ± 17.3 AZs, n=13, F (5, 60) = 62.05, p=0.0068, Figure 6A), AZ number contributed solely by the MN1-Ib input was not significantly altered whether Is was absent (285.8 ± 29.7 Ib AZs, n=10) or present (329.1 ± 15.8 Ib AZs, n=13, F (5, 60) = 62.05, p=0.66, Figure 6A). Similarly, no change in synaptic bouton number in MN1-Ib motoneurons was observed whether MNIs was absent (18.0 ± 4.3 Ib boutons, n=8) or present (19.1 ± 4.4 Ib boutons, n=16, F (5, 66) = 60.73, p=0.99, Figure 6B). We conclude that synaptic growth of MN1-Ib is not altered when muscle M1 lacks Is innervation.

**Figure 6.**
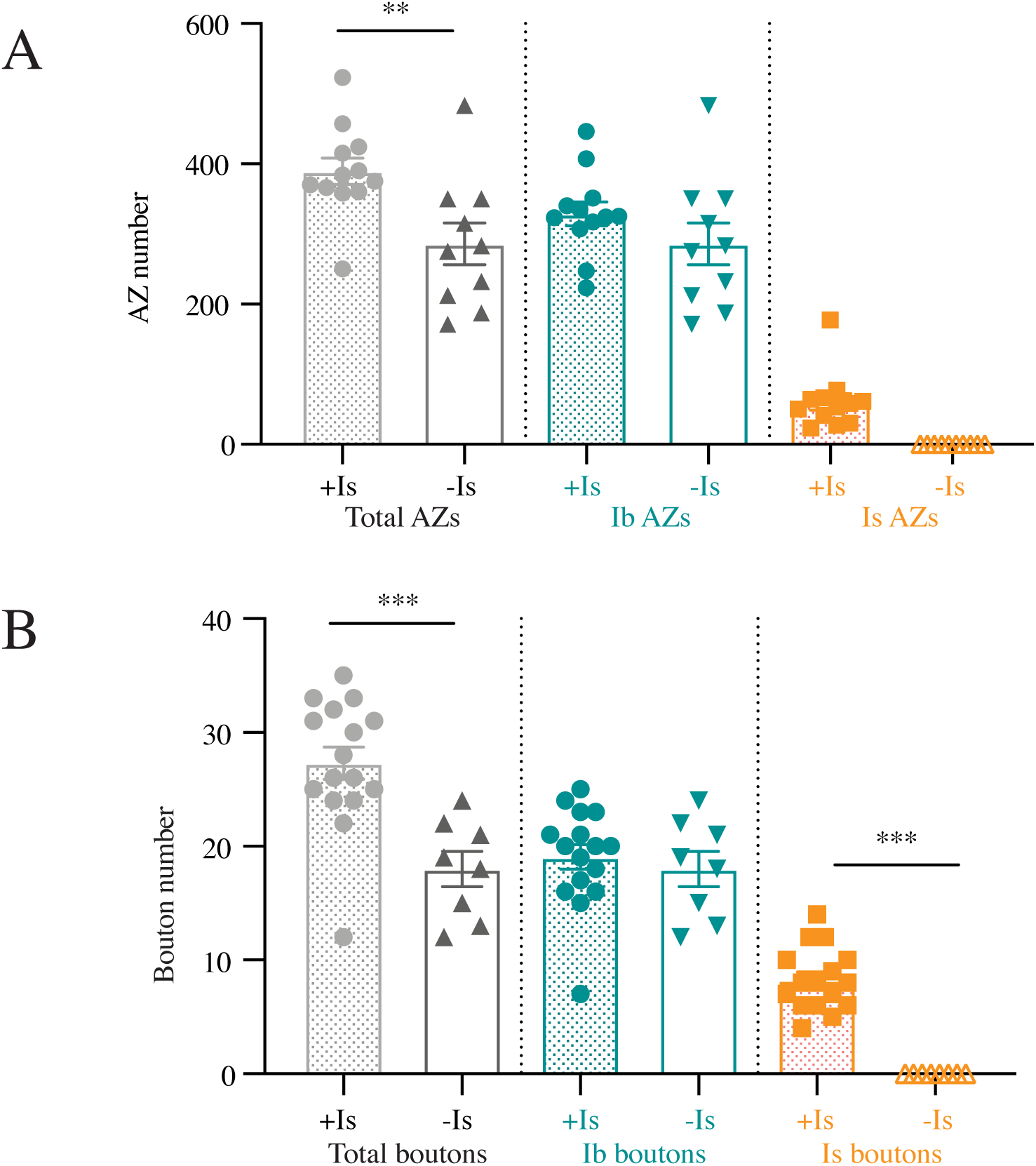
Lack of structural synaptic changes in MN1-Ib when Is innervation is absent. ***A***, Quantification of MN1-Ib and MNIs AZ number following immunolabeling for BRP in control *w*^*118*^ 3^rd^ instar larval M1 muscles in segment A3. Total AZ number when both inputs are present (+Is) or when Is innervation is absent (-Is) is shown. AZ number specifically for MN1-Ib (teal) or MNIs (orange) is also shown when both inputs are present (+Is) or when Is innervation is absent (-Is). ***B***, Quantification of MN1-Ib and MNIs synaptic bouton number following immunolabeling for HRP in *w*^*118*^ 3^rd^ instar larval M1 muscles in segment A3. Total bouton number when both inputs are present (+Is) or when Is innervation is absent (-Is) is shown. Bouton number specifically for MN1-Ib (teal) or MNIs (orange) is also shown when both inputs are present (+Is) or when Is innervation is absent (-Is). Each data point represents quantification from a single larva. Statistical significance was determined using ANOVA. Data is shown as mean ± SEM; ** = p<0.01.

### Ablation of MNIs triggers increased evoked release from the remaining MN1-Ib input

Although no structural compensation was observed in MN1-Ib when MNIs was absent, functional changes in neurotransmitter release could occur in the absence of increased release sites. In addition, a mismatch in neuronal activity between inputs during development could result in unique forms of plasticity compared to when a motoneuron was missing. To generate an input imbalance, MN1-Ib or MNIs GAL4 drivers were used to express several well-characterized UAS-transgenes that alter neuronal activity (Simpson, 2009; Venken et al., 2011; Yoshihara and Ito, 2012; Pauls et al., 2015). To decrease neurotransmitter release and synaptic output, a transgene encoding tetanus toxin light chain (UAS-TeTXLC) was expressed to cleave the v-SNARE n-Synaptobrevin and eliminate evoked synaptic transmission (Sweeney et al., 1995). A transgene encoding a bacterial voltage-gated Na^+^ channel (UAS-NaChBac) that enhances depolarization was used to constitutively increase neuronal excitability (Nitabach et al., 2006). To compare the effects of reduced or enhanced activity with the complete absence of each input, Reaper expression (UAS-RPR) was used to ablate MN1-Ib or MNIs. None of the manipulations altered M1 muscle surface area (F (10, 175) = 2.129, p=0.66, n=6 to 29 per genotype), indicating muscle growth is not affected by ablation or activity changes in MN1-Ib or MNIs motoneurons (Figure 7–1). For all experimental manipulations, the Ib motoneuron was labeled with MN1-Ib LexA, LexAop-GFP in the background for unambiguous identification of the two inputs following immunostaining with the pan-neuronal marker anti-HRP (Ib and Is) and anti-GFP (Ib only). Synaptic development, synaptic function and muscle contraction force were analyzed in controls and compared to Ib and Is motoneurons expressing the transgenes with their respective GAL4 driver.

**Figure 7.**
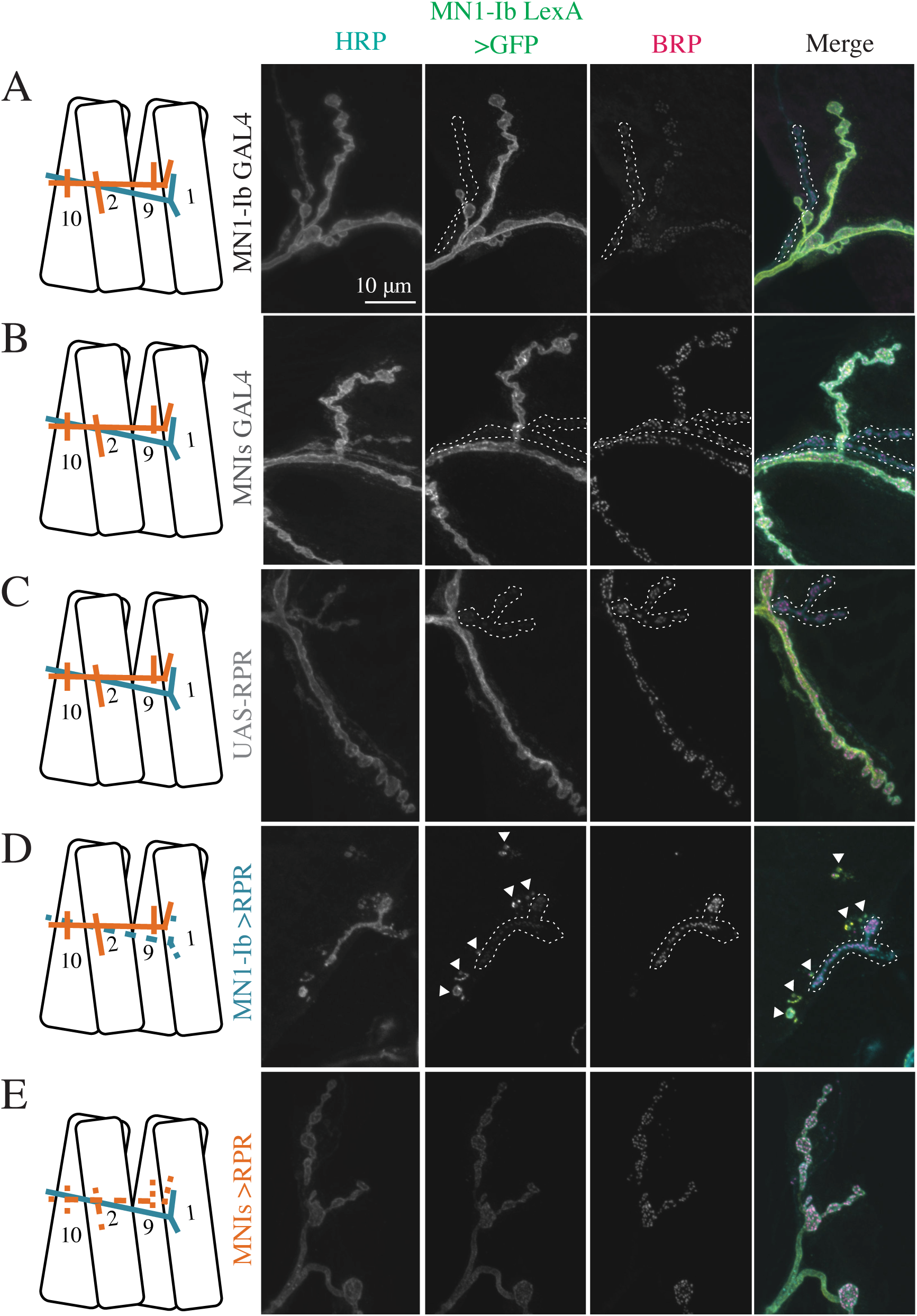
Morphological consequences of ablation of MN1-Ib or MNIs. Representative confocal images of 3^rd^ instar larval M1 NMJs at segment A3 following immunolabeling with anti-HRP, anti-GFP and anti-BRP in the following genotypes: ***A***, MN1-Ib GAL4 control; ***B***, MNIs GAL4 control; ***C***, UAS-RPR control; ***D***, MN1-Ib GAL4>UAS-RPR; ***E***, MNIs GAL4>UAS-RPR. MN1-Ib LexA>LexAop2-CD8-GFP was present in each genetic background to allow unambiguous identification of the Ib terminal. Diagrams of the experimental manipulation is shown on the left, with MN1-Ib (teal) and MNIs (orange) labeled. The merged image is shown on the right. The white dashed line highlights the MNIs terminal in the final 3 panels for each manipulation except for ***E***, where Is is absent following ablation. Arrowheads in ***D*** depict GFP-positive debris near M1 secondary to death and fragmentation of MN1-Ib following Reaper expression. Scale bar = 10 *µ*m for all panels.

**Figure 7-1.**
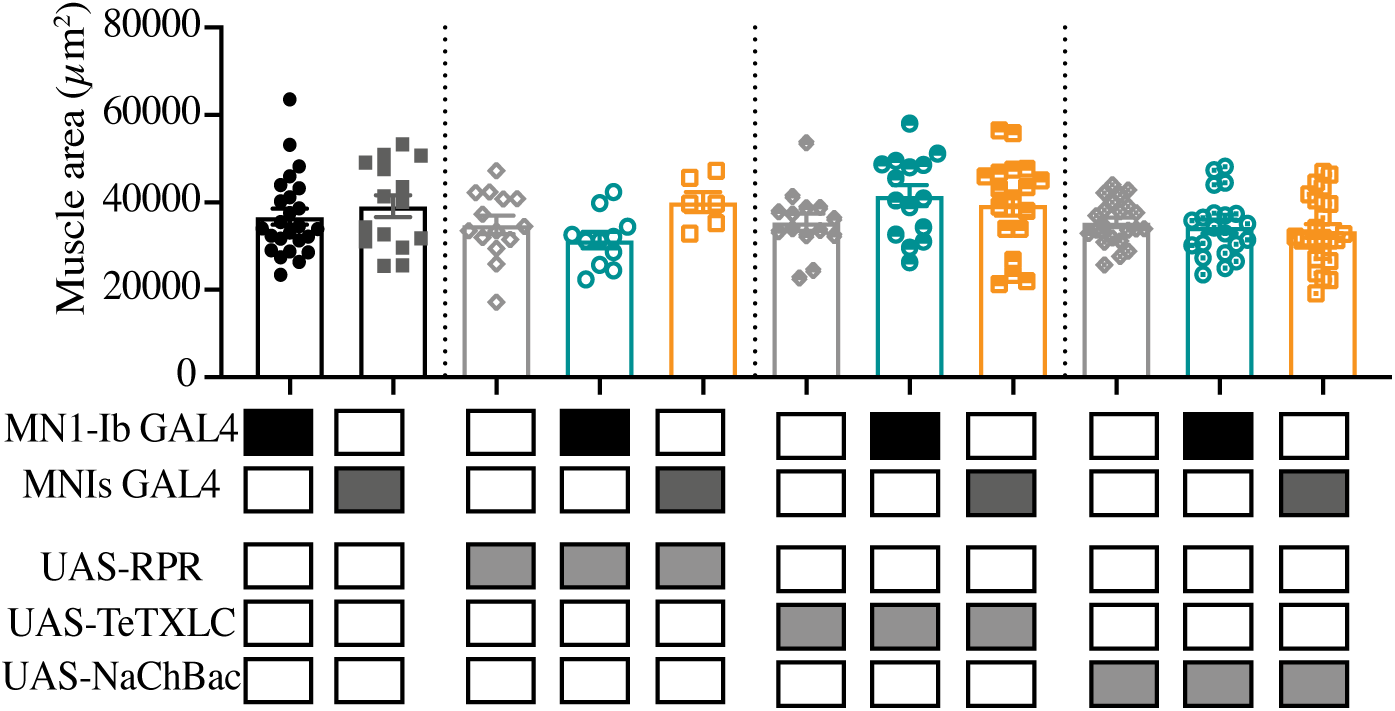
M1 muscle size is not altered by ablation or activity changes of MN1-Ib or MNIs motoneurons. Shaded boxes under each bar indicate the genotypes for each group, with control GAL4 driver lines alone (MN1-Ib, MNIs), control UAS transgenes alone (UAS-RPR, UAS-TeTXLC, UAS-NaChBac) and experimental crosses of MN1-Ib GAL4 (teal) or MNIs GAL4 (orange) to each transgene. Each data point represents quantification of segment A3 M1 surface area from a single 3^rd^ instar larvae. Statistical significance was determined using ANOVA. No statistical difference was found across genotypes. Data is shown as mean ± SEM.

We first examined if ablation of each motoneuron subclass altered synaptic structure of the remaining input. Expression of UAS-RPR with either MN1-Ib GAL4 or MNIs GAL4 resulted in elimination of the respective motoneuron compared to controls (Figure 7A-E). Membrane fragments immuno-positive for GFP from the MN1-Ib LexA, LexAop-GFP labeling were often observed near M1 in MN1-Ib GAL4>UAS-RPR larvae (Figure 7D), suggesting cell death and membrane fragmentation occurred after the initial stages of axonal pathfinding. Genetic ablation of the phasic Is motoneuron in MNIs GAL4>UAS-RPR larvae caused a mild decrease in total AZ and synaptic bouton number due to loss of the Is input (n=14 to 15, F(10, 183) = 25.17, p= 0.03, Figure 8A, B). However, there was no change in MN1-Ib AZ (F (10, 183) = 52.42, p=0.999, Figure 9A) or bouton number (F (10, 185) = 29.99, p=0.999, Figure 9B) following MNIs ablation, similar to conditions where MNIs naturally failed to innervate M1 in control larvae.

**Figure 8.**
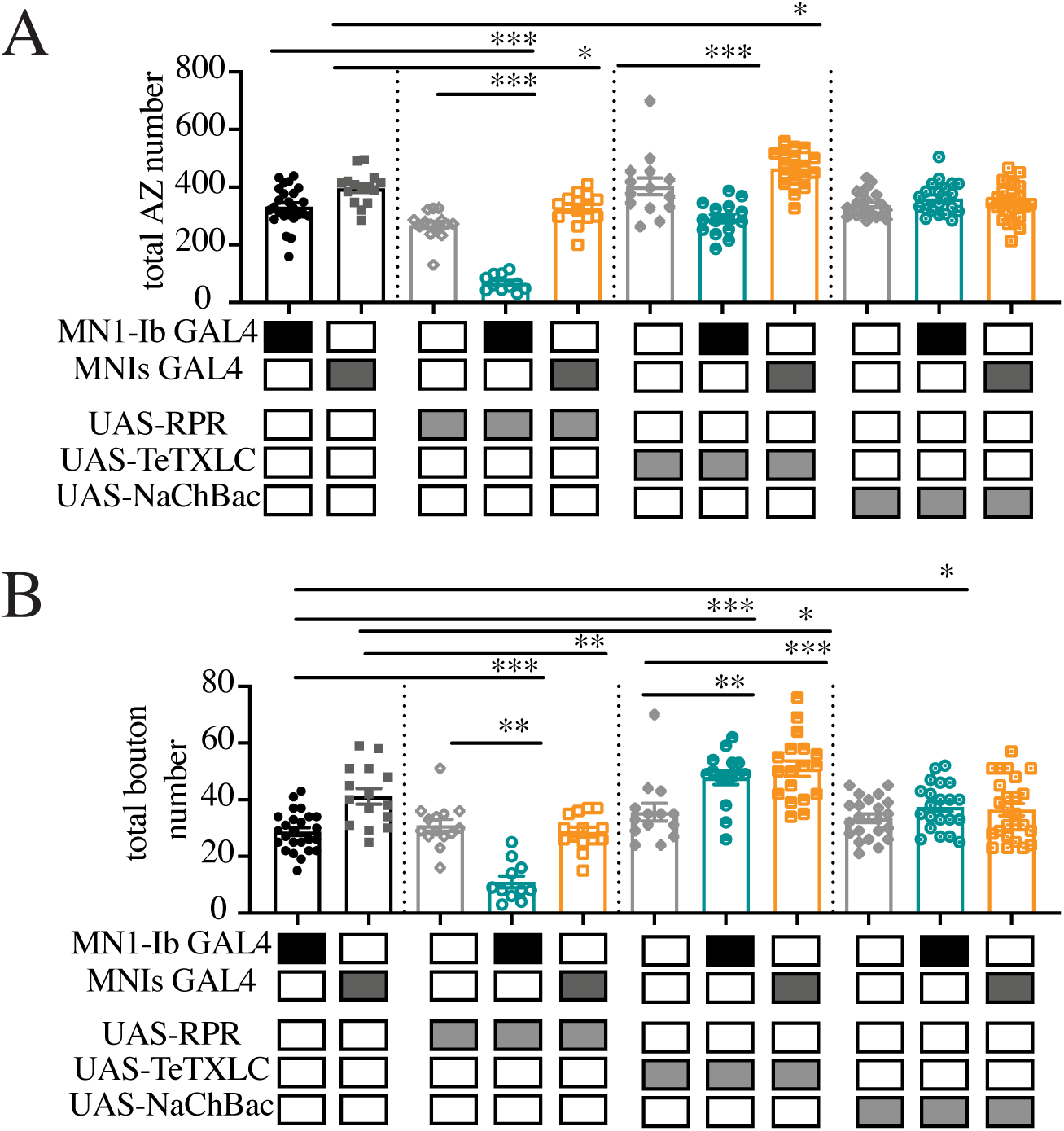
Quantification of total AZ and bouton number follow ablation or activity changes of MN1-Ib or MNIs. ***A***, Quantification of combined MN1-Ib and MNIs AZ number following immunolabeling for BRP in 3^rd^ instar larval M1 muscles in segment A3 of the indicated genotypes. ***B***, Quantification of combined MN1-Ib and MNIs synaptic bouton number following immunolabeling for HRP in 3^rd^ instar larval M1 muscles in segment A3 of the indicated genotypes. Shaded boxes under each bar indicate the genotypes for each group, with control GAL4 driver lines alone (MN1-Ib, MNIs), control UAS transgenes alone (UAS-RPR, UAS-TeTXLC, UAS-NaChBac) and experimental crosses of MN1-Ib GAL4 (teal) or MNIs GAL4 (orange) to each transgene. Each data point represents quantification from segment A3 M1 from a single 3^rd^ instar larvae. Statistical significance was determined using ANOVA. Data is shown as mean ± SEM; * = p<0.05, ** = p<0.01, *** = p<0.001.

**Figure 9.**
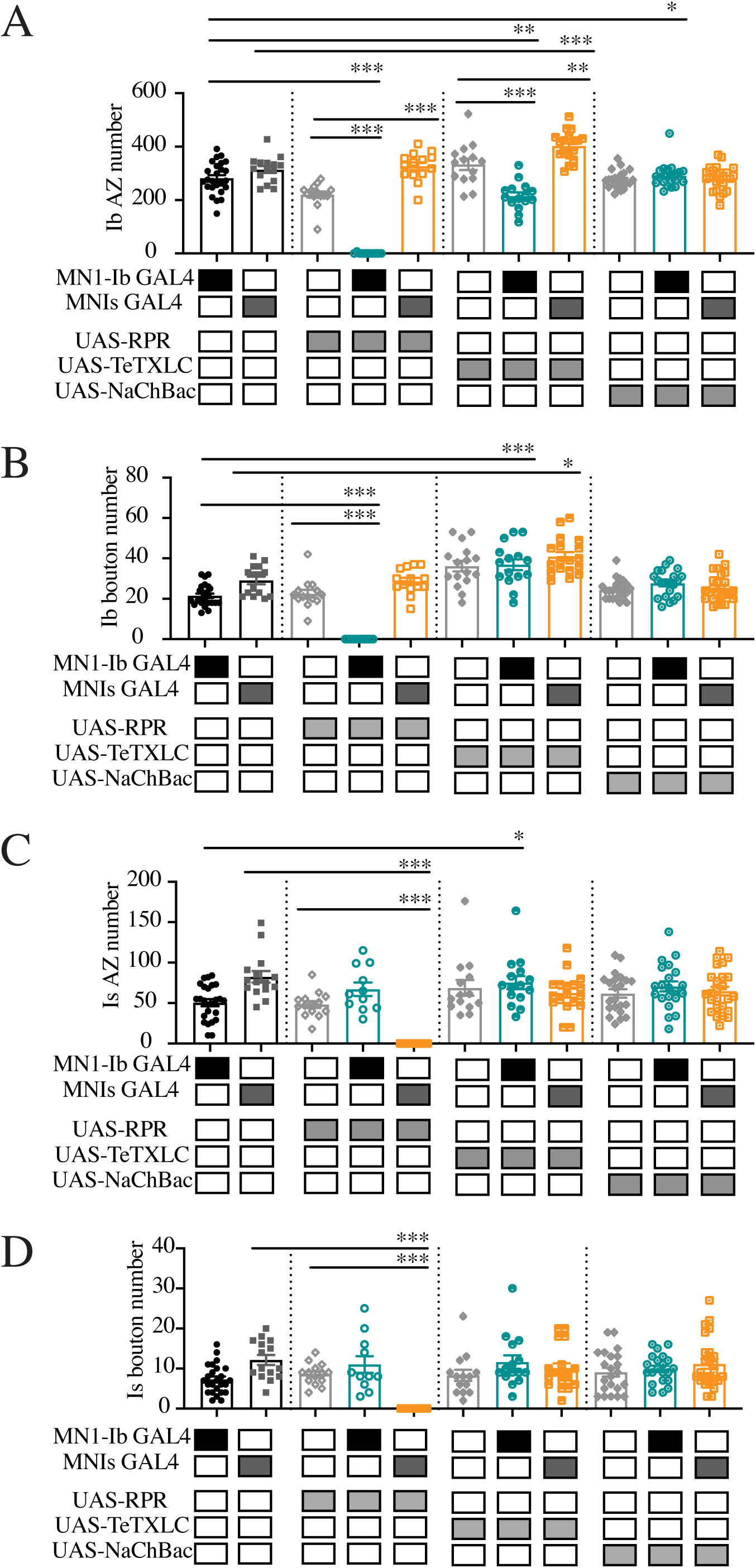
Quantification of MN1-Ib or MNIs AZ and bouton number follow ablation or activity changes. ***A***, Quantification of MN1-Ib AZ number following immunolabeling for BRP in 3^rd^ instar larval M1 muscles in segment A3 of the indicated genotypes. ***B***, Quantification of combined MN1-Ib synaptic bouton number following immunolabeling for HRP in 3^rd^ instar larval M1 muscles in segment A3 of the indicated genotypes. ***C***, Quantification of MNIs AZ number following immunolabeling for BRP in 3^rd^ instar larval M1 muscles in segment A3 of the indicated genotypes. ***D***, Quantification of MNIs synaptic bouton number following immunolabeling for HRP in 3^rd^ instar larval M1 muscles in segment A3 of the indicated genotypes. Shaded boxes under each bar indicate the genotypes for each group, with control GAL4 driver lines alone (MN1-Ib, MNIs), control UAS transgenes alone (UAS-RPR, UAS-TeTXLC, UAS-NaChBac) and experimental crosses of MN1-Ib GAL4 (teal) or MNIs GAL4 (orange) to each transgene. Each data point represents quantification from segment A3 M1 from a single 3^rd^ instar larva. Statistical significance was determined using ANOVA. Data is shown as mean ± SEM; * = p<0.05, ** = p<0.01, *** = p<0.001.

To determine if MN1-Ib altered its functional properties without changes in the number of release sites, electrophysiology was performed at 3^rd^ instar M1 muscles to measure Ib evoked release when Is innervation was present in control versus when Is was ablated with UAS-RPR. Although no structural changes were identified at MN1-Ib NMJs, a functional change in the output of the tonic motoneuron was observed. The evoked EJP response triggered by MN1-Ib activation was increased 24% at M1 muscles when MNIs input was ablated (15.4 ± 1.0 mV, n=11) versus when MNIs was present (11.8 ± 1.6 mV, n=22, F (10, 58) = 5.30, p=0.05, Figure 10). This compensation did not result in complete recovery of the evoked output observed when both inputs were present (control Is-GAL4: 25.9 ± 1.5 mV, n=10). In addition, contractile force was still decreased following MNIs ablation as previously noted (Figure 4E-G, Figure 10–1A, C). These data suggest the muscle is capable of detecting loss of the Is input and increasing synaptic output from the remaining Ib motoneuron, resulting in a partial rescue of muscle excitability.

**Figure 10.**
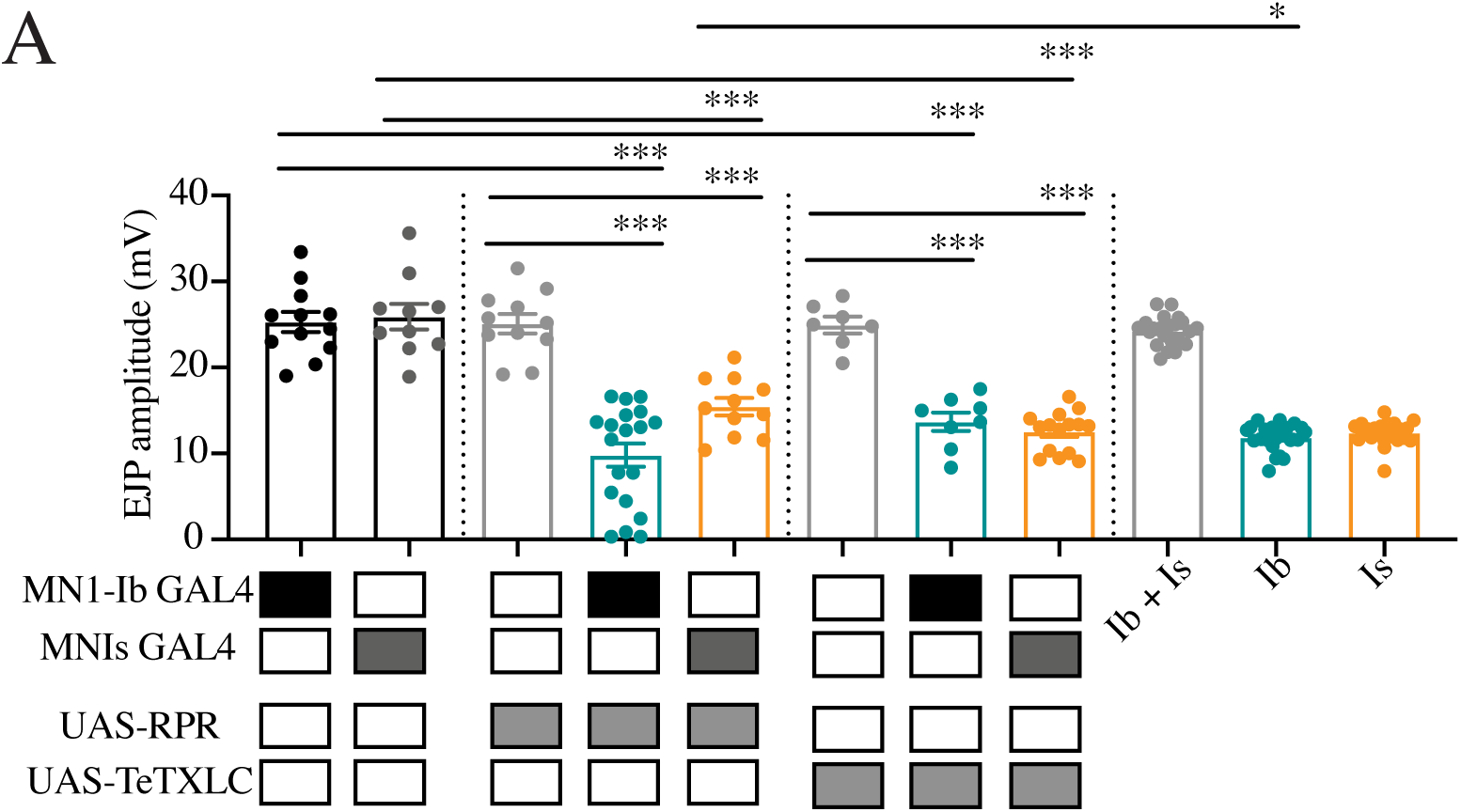
Electrophysiological analysis of synaptic function at M1 following ablation or silencing of MN1-Ib or MNIs. ***A***, EJP amplitudes recorded from 3^rd^ instar larval M1 muscles in segment A3 of the indicated genotypes. Each data point is the average of at least 20 EJPs recorded from each larva. Shaded boxes under each bar indicate the genotypes for each group, with control GAL4 driver lines alone (MN1-Ib, MNIs), control UAS transgenes alone (UAS-RPR, UAS-TeTXLC) and experimental crosses of MN1-Ib GAL4 (teal) or MNIs GAL4 (orange) to each transgene. The final 3 bars on the right show results from dual intracellular recordings in controls using the minimal stimulation protocol where both MN1-Ib or MNIs motoneurons were active (Ib+Is), or MN1-Ib (Ib) or MNIs (Is) were independently isolated. Statistical significance was determined using ANOVA. Data is shown as mean ± SEM; * = p<0.05, *** = p<0.001.

**Figure 10–1.**
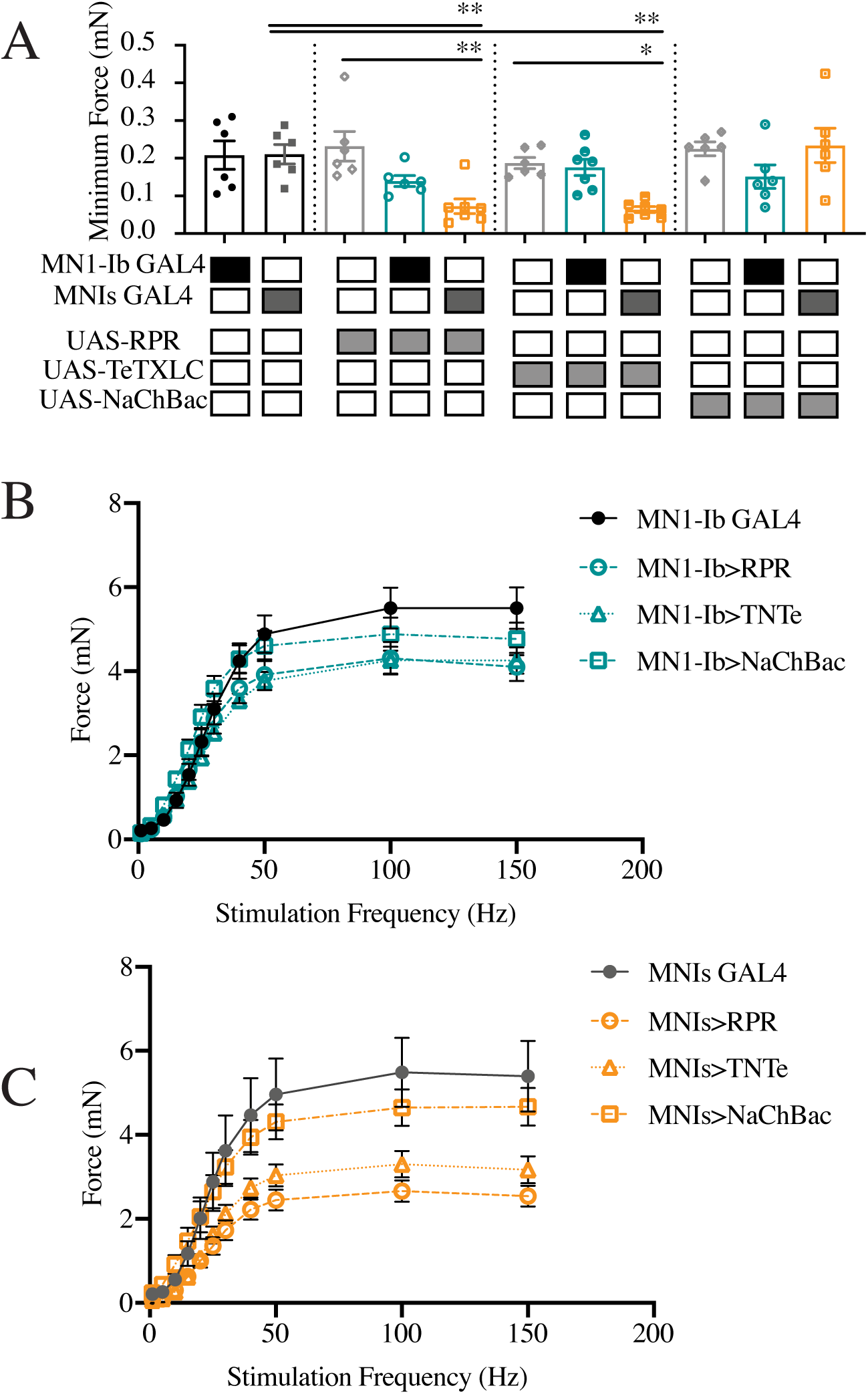
Force recordings from muscles following ablation or activity changes of MN1-Ib or MNIs. ***A***, Maximal contraction force elicited by 150 Hz stimulation in 3^rd^ instar larvae of the indicated genotypes. Six replicate contractions per genotype were generated at each stimulation frequency per recording. Shaded boxes under each bar indicate the genotypes for each group, with control GAL4 driver lines alone (MN1-Ib, MNIs), control UAS transgenes alone (UAS-RPR, UAS-TeTXLC, UAS-NaChBac) and experimental crosses of MN1-Ib GAL4 (teal) or MNIs GAL4 (orange) to each transgene. Statistical significance was determined using ANOVA. Data is shown as mean ± SEM; * = p<0.05, ** = p<0.01. ***B***, Force-frequency curves for MN1-Ib GAL4 controls and the indicated experimental genotypes. Data points represent 6 replicate contractions elicited at each frequency from 6-7 3^rd^ instar larvae. ***C***, Force-frequency curves for MN1-Is GAL4 controls and the indicated experimental genotypes. Data points represent 6 replicate contractions elicited at each frequency from 6-7 3^rd^ instar larvae.

To examine if phasic motoneurons displayed similar functional compensation, genetic ablation of the tonic Ib motoneuron was performed using MN1-Ib GAL4; UAS-RPR. As previously described, M1 occasionally lacked MNIs input due to natural variation in controls. As such, MN1-Ib ablation resulted in M1 having no synaptic innervation (42%) or only MNIs innervation (58%). Larvae with Is innervation at M1 following ablation of the Ib motoneuron displayed a dramatic decrease in the number of total AZs (control: 332.1 ± 13.5 AZs, n=25; Ib>RPR: 67.7 ± 8.4 AZs, n=11, F (10, 183) = 35.17, p<0.0001, Figure 8A) and synaptic boutons (control: 28.8 ± 1.4, n=25; Ib>RPR: 11.0 ± 2.1, n=11, F (10, 183) = 19.47, p<0.0001, Figure 8B), given the larger number of synapses normally contributed by the MN1-Ib input. However, loss of MN1-Ib did not trigger changes in the number of Is AZs (Figure 9C) or Is synaptic boutons (Figure 9D) compared to Is innervation when MN1-Ib was present. In contrast to the functional increase in evoked release in tonic Ib neurons following ablation of Is, loss of MN1-Ib did not trigger a compensatory increase in evoked output from the remaining MNIs motoneuron (Figure 10). We conclude that compensatory structural or functional changes do not occur in the phasic Is input following loss of the tonic Ib motoneuron at M1. In contrast, loss of Is results in functional changes in the co-innervating Ib input that partially compensate for the reduced evoked response.

### Imbalances in MN1-Ib or MNIs neuronal activity reveal structural plasticity of tonic Ib inputs

To examine the consequences of activity perturbations between Ib and Is motoneurons at M1 NMJs during development, manipulations were performed to increase or decrease synaptic output of one of the two neurons. Expression of UAS-TeTXLC in either the Ib or Is input blocked evoked synaptic transmission from the affected motoneuron, reducing EJP amplitude recorded physiologically to the level observed when only the Ib or Is motoneuron were recruited during minimal stimulation (Figure 10). Silencing the Is motoneuron in MNIs GAL4; UAS-TeTXLC larvae resulted in structural changes at the NMJ, with an increase in the total number of AZs (control: 396.3 ± 14.6 AZs, n=15; MNIs>TeTXLC: 465.0 ± 14.7 AZs, n=18, F (10, 183) = 35.17, p=0.0187, Figure 8A) and synaptic boutons (control: 41.2 ± 2.7, n=15; Is>TeTXLC: 50.9 ± 2.7, n=18, F (10, 183) = 19.47, p=0.028, Figure 8B). This enhanced synaptic growth was due to increases occurring in the co-innervating MN1-Ib input (Figure 9A, B), with no changes observed in the affected MNIs (Figure 9C, D). In particular, MN1-Ib displayed a large increase in AZ number when Is was silenced compared to when Is was present or ablated (Is present: 314 ± 12.9, n=15; Is ablated: 326.6 ± 14.4, n=14; Is silenced: 402.7 ± 13.0, n=18, F (10, 183) = 35.17, p=0.0187). Although the number of release sites increased in MN1-Ib following silencing of MNIs (MNIs>TeTXLC), electrophysiology (Figure 10) and contraction assays (Figure 10 – supplemental figure 1A, C) indicated these structural changes were insufficient to induce increased excitability or contractility of the muscle. We conclude that the complete absence of the phasic Is input leads to a functional increase in release from the co-innervating tonic Ib input without a change in the number of release sites. In contrast, when Is is present but functionally silent, the tonic Ib input displays a distinct response with a structural change that includes more release sites, but the overall functional output of the motoneuron remains unaltered.

We next examined the consequences of silencing MN1-Ib activity with TeTXLC. Similar to when MN1-Ib was ablated, the co-innervating MNIs did not display structural (Figure 9C, D) or functional compensation (Figure 10, Figure 10-1A, B), indicating the phasic motoneuron is less capable of compensatory synaptic plasticity when the co-innervating tonic motoneuron is ablated or silenced. In contrast to the lack of change in the phasic Is input, silencing MN1-Ib (Ib>TeTXLC) triggered several structural changes to its own morphology. First, a striking reduction in AZ number was found, with a 30% decrease in release sites in MN1-Ib motoneurons lacking evoked transmission (UAS-TeTXLC: 333.6 ± 21.2 AZs, n=14; MN1-Ib GAL4: 281.6 ± 11.9 AZs, n=25; MN1-Ib>TeTXLC: 215.3 ± 13.6 AZs, n=15, F (10, 183) = 52.42, p=0.001, Figure 9A). Second, there was a change in the anatomy of the MN1-Ib axon at the NMJ, with the appearance of synaptic filopodial-like protrusions (Figure 11A-C). This phenotype was never observed in controls (average protrusions per MN1-Ib NMJ: UAS-TeTXLC: 0, n=14; MN1-Ib GAL4: 0, n=25; MN1-Ib>TeTXLC: 7.2 ± 2.4, n=16, F (4, 83) = 9.921, p=0.0001, Figure 11D). Similar filopodial-like protrusions were described previously at early 1^st^ instar NMJs during the initial stages of synapse formation in wildtype animals, but never at mature 3^rd^ instar NMJs (Akbergenova et al., 2018). Such protrusions were not observed in silenced Is motoneurons or in Ib motoneurons following Is silencing (Figure 11A-D), indicating MN1-Ib and MNIs react differently to changes in their own activity.

**Figure 11.**
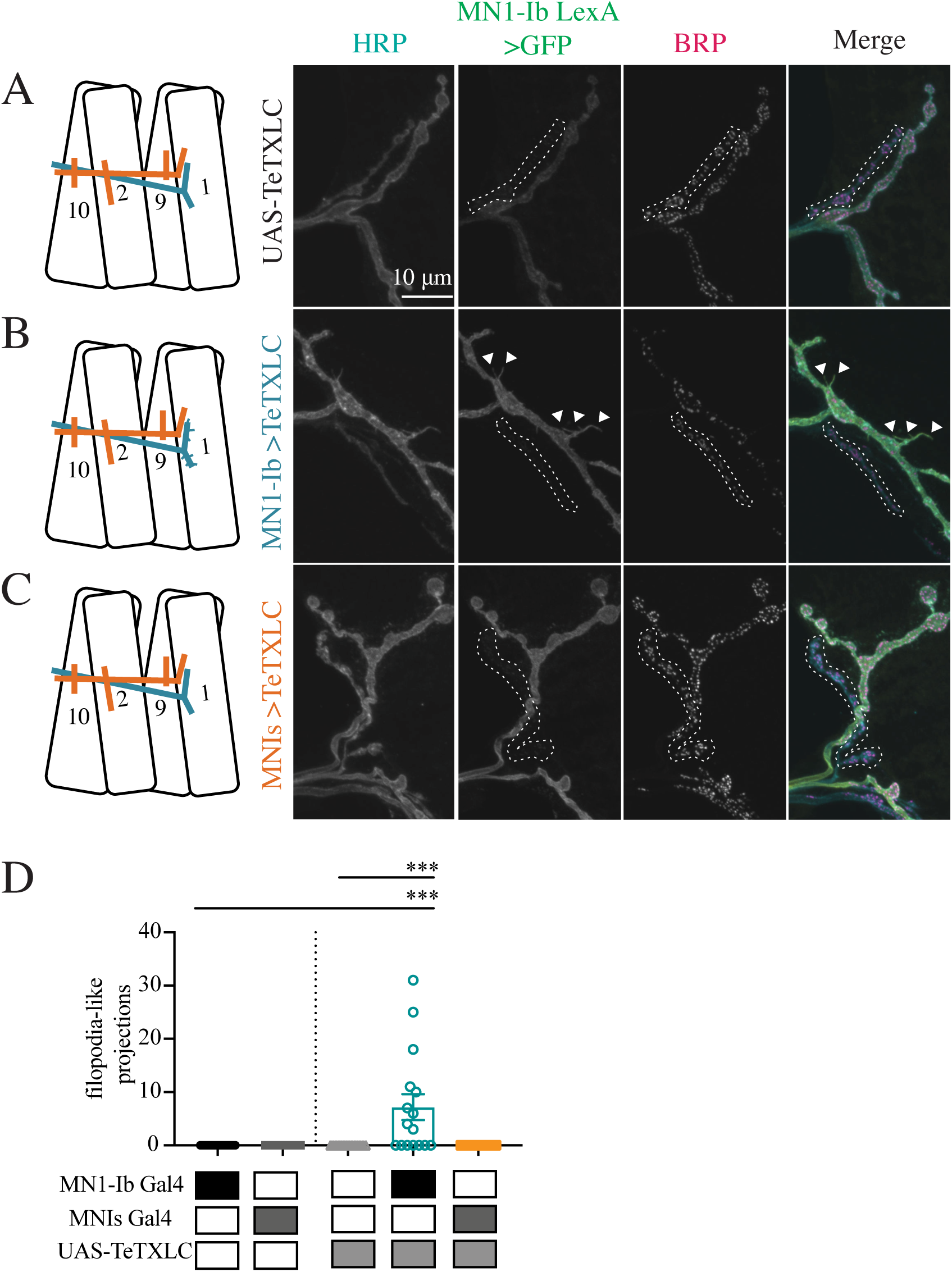
Morphological consequences of silencing of MN1-Ib or MNIs. Representative confocal images of 3^rd^ instar larval M1 NMJs at segment A3 following immunolabeling with anti-HRP, anti-GFP and anti-BRP in the following genotypes: ***A***, UAS-TeTXLC control; ***B***, MN1-Ib GAL4>UAS-TeTXLC; ***C***, MNIs GAL4>UAS-TeTXLC. MN1-Ib LexA>LexAop2-CD8-GFP was present in each genetic background to allow unambiguous identification of the Ib terminal. Diagrams of the experimental manipulation is shown on the left, with MN1-Ib (teal) and MNIs (orange) labeled. The merged image is shown on the right. The white dashed line highlights the MNIs terminal in the final 3 panels for each manipulation. Arrowheads in ***B*** depict GFP-positive filopodial-like projections from MN1-Ib following tetanus toxin expression. Scale bar = 10 *µ*m for all panels. ***D***, Quantification of filopodial-like projections in controls and following UAS-TeTXLC expression with MN1-Ib or MNIs GAL4. Each data point represents quantification from segment A3 M1 from a single 3^rd^ instar larvae. Statistical significance was determined using ANOVA. Data is shown as mean ± SEM; *** = p<0.001.

A final morphological change at silenced MN1-Ib NMJs was a decrease in postsynaptic SSR membrane revealed by anti-DLG staining (Figure 12A-D). SSR volume compared to presynaptic NMJ volume (anti-HRP staining) was reduced by 50% on average at MN1-Ib>TeTXLC NMJs compared to controls (F (2, 54) = 11.70, p=0.0079, Figure 12D). Together, the reduced AZ number, decreased muscle SSR volume and increased synaptic filopodial-like protrusions suggest silenced MN1-Ib motoneurons maintain an immature state with reduced AZ formation and a failure to properly induce normal postsynaptic specializations. These defects are not observed at silenced Is phasic synapses, indicating the phasic Is motoneuron class is less sensitive to activity changes and any potential compensatory responses triggered from the muscle.

**Figure 12.**
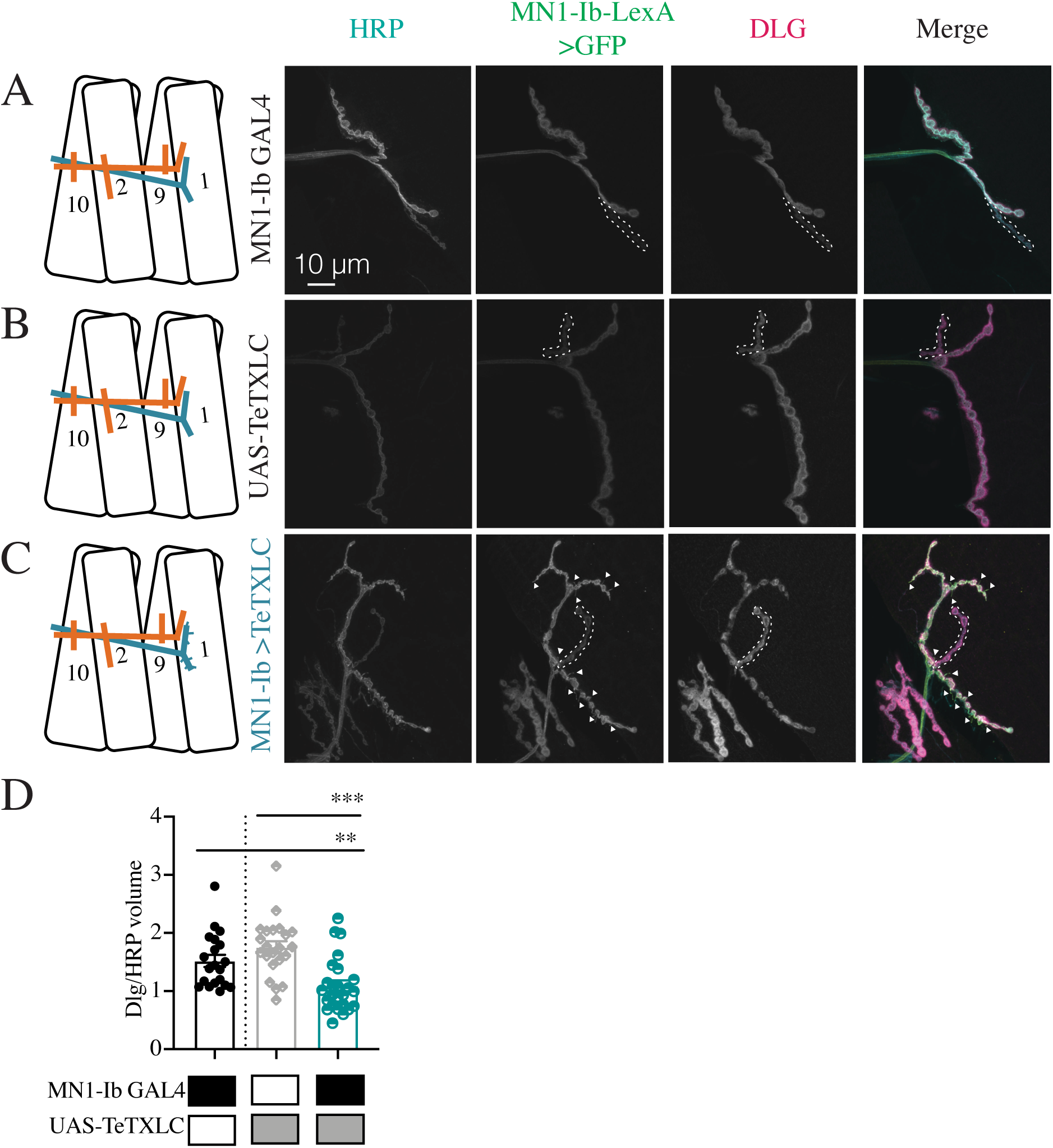
Reduced postsynaptic SSR volume following silencing of MN1-Ib. Representative confocal images of 3^rd^ instar larval M1 NMJs at segment A3 following immunolabeling with anti-HRP, anti-GFP and anti-DLG in the following genotypes: ***A***, MN1-Ib GAL4 control; ***B***, UAS-TeTXLC control; ***C***, MN1-Ib GAL4>UAS-TeTXLC. MN1-Ib LexA>LexAop2-CD8-GFP was present in each genetic background to allow unambiguous identification of the Ib terminal. Diagrams of the experimental manipulation is shown on the left, with MN1-Ib (teal) and MNIs (orange) labeled. The merged image is shown on the right. The white dashed line highlights the MNIs terminal in the final 3 panels for each manipulation. Arrowheads in ***C*** depict GFP-positive filopodial-like projections from MN1-Ib following tetanus toxin expression. Scale bar = 10 *µ*m for all panels. ***D***, Quantification of DLG to HRP volume in MN1-Ib in controls and following UAS-TeTXLC expression with MN1-Ib GAL4. Each data point represents quantification from segment A3 M1 from a single 3^rd^ instar larvae. Statistical significance was determined using ANOVA. Data is shown as mean ± SEM; ** = p<0.01, *** = p<0.001.

To determine if enhanced activity of either of the two motoneuron subclasses would induce structural or functional changes, the NaChBac depolarizing Na^+^ channel was expressed in either MN1-Ib or MNIs (Figure 13A-C). NaChBaC expression has been previously demonstrated to enhance membrane depolarization by increasing Na^+^ conductance (Pauls et al., 2015). Consistent with enhanced excitability and increased burst spiking in affected motoneurons, trains of EJPs in response to a single stimulus were often recorded from M1 in larvae expressing NaChBac (Figure 13D). Although expression of the channel enhanced excitability, it did not result in structural (Figure 8, Figure 9) or functional (Figure 13E, Figure 10-1) changes in synaptic properties of MN1-Ib>NaChBac or MNIs>NaChBac larvae. No alterations of the Ib or Is input were observed in either condition. Similarly, increased activity in either motoneuron class did not trigger any obvious structural competition between the inputs. We conclude that NMJ plasticity is more sensitive to manipulations that reduce presynaptic release versus those that enhance membrane excitability, and that these changes preferentially manifest within the tonic Ib subclass of motoneurons.

**Figure 13.**
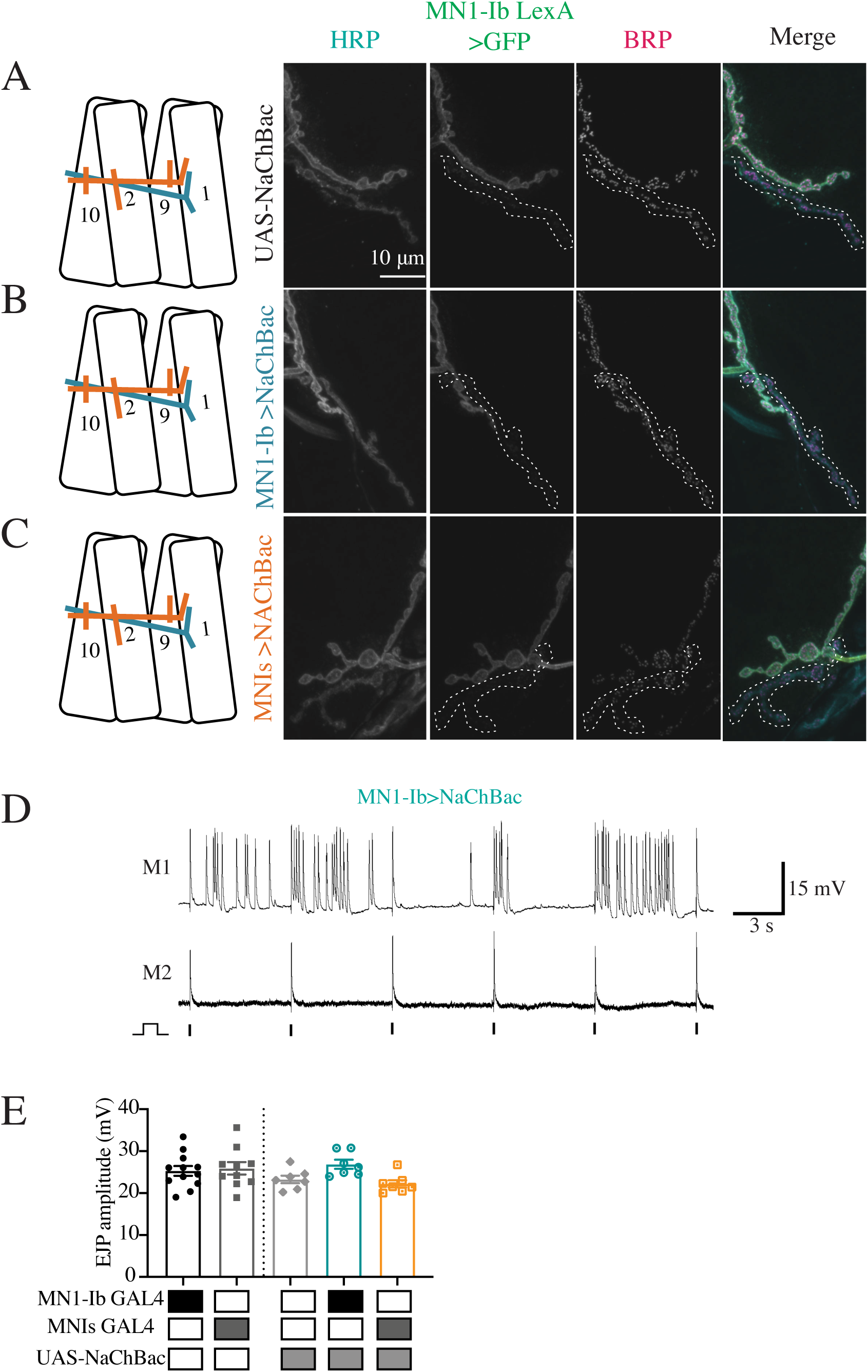
Chronic increases in MN1-Ib or MNIs activity do not impact NMJ morphology or synaptic release. Representative confocal images of 3^rd^ instar larval M1 NMJs at segment A3 following immunolabeling with anti-HRP, anti-GFP and anti-BRP in the following genotypes: ***A***, UAS-NaChBac control; ***B***, MN1-Ib GAL4>UAS-NaChBac; ***C***, MNIs GAL4>UAS-NaChBac. MN1-Ib LexA>LexAop2-CD8-GFP was present in each genetic background to allow unambiguous identification of the Ib terminal. Diagrams of the experimental manipulation is shown on the left, with MN1-Ib (teal) and MNIs (orange) labeled. The merged image is shown on the right. The white dashed line highlights the MNIs terminal in the final 3 panels for each manipulation. Scale bar = 10 *µ*m for all panels. ***D***, Representative dual intracellular recordings from M1 and M2 in MN1-Ib>NaChBac 3^rd^ instar larvae during 0.2 Hz stimulation. Note the train of EJPs following a single stimulus at M1 compared to M2. Vertical lines below the M2 recordings indicate timing of nerve stimulation. ***E***, EJP amplitudes recorded from 3^rd^ instar larval M1 muscles in segment A3 of the indicated genotypes. Each data point is the average of at least 20 EJPs recorded from each larva. Statistical significance was determined using ANOVA. No statistical difference was found across genotypes. Data is shown as mean ± SEM.

## Discussion

To characterize how changes in the presence or activity of tonic versus phasic motoneurons alter NMJ development and function in Drosophila, we identified GAL4 drivers specific for the Ib and Is neuronal subclasses that innervate M1 and used them to alter the balance of input to the muscle. Our data indicate Ib and Is glutamatergic motoneurons largely form independent inputs that make similar contributions to muscle excitability and contractile force. The tonic Ib subclass was capable of structural or functional changes following manipulations that altered their output or that of the co-innervating phasic Is motoneuron (Figure 14A-D). These changes were only observed during conditions when neuronal activity was decreased or when the Is input was ablated. Functional increases in evoked release without enhanced synapse number were observed in Ib motoneurons following ablation of Is (Figure 14B). In contrast, morphological changes that increased AZ number without enhancing evoked release occurred when Is synaptic output was blocked with tetanus toxin (Figure 14C). While Ib motoneurons were capable of several forms of plastic change following reduced input to the muscle, the phasic Is motoneurons were insensitive to manipulations of their own activity or that of the tonic Ib input. Unlike plasticity observed in Ib neurons following reduction in synaptic drive to the muscle, enhancing excitability of either the Ib or Is input was ineffective at triggering changes in either motoneuron class. These data indicate reductions in activity from either input trigger a structural or functional change primarily from the tonic Ib motoneuron subclass.

**Figure 14.**
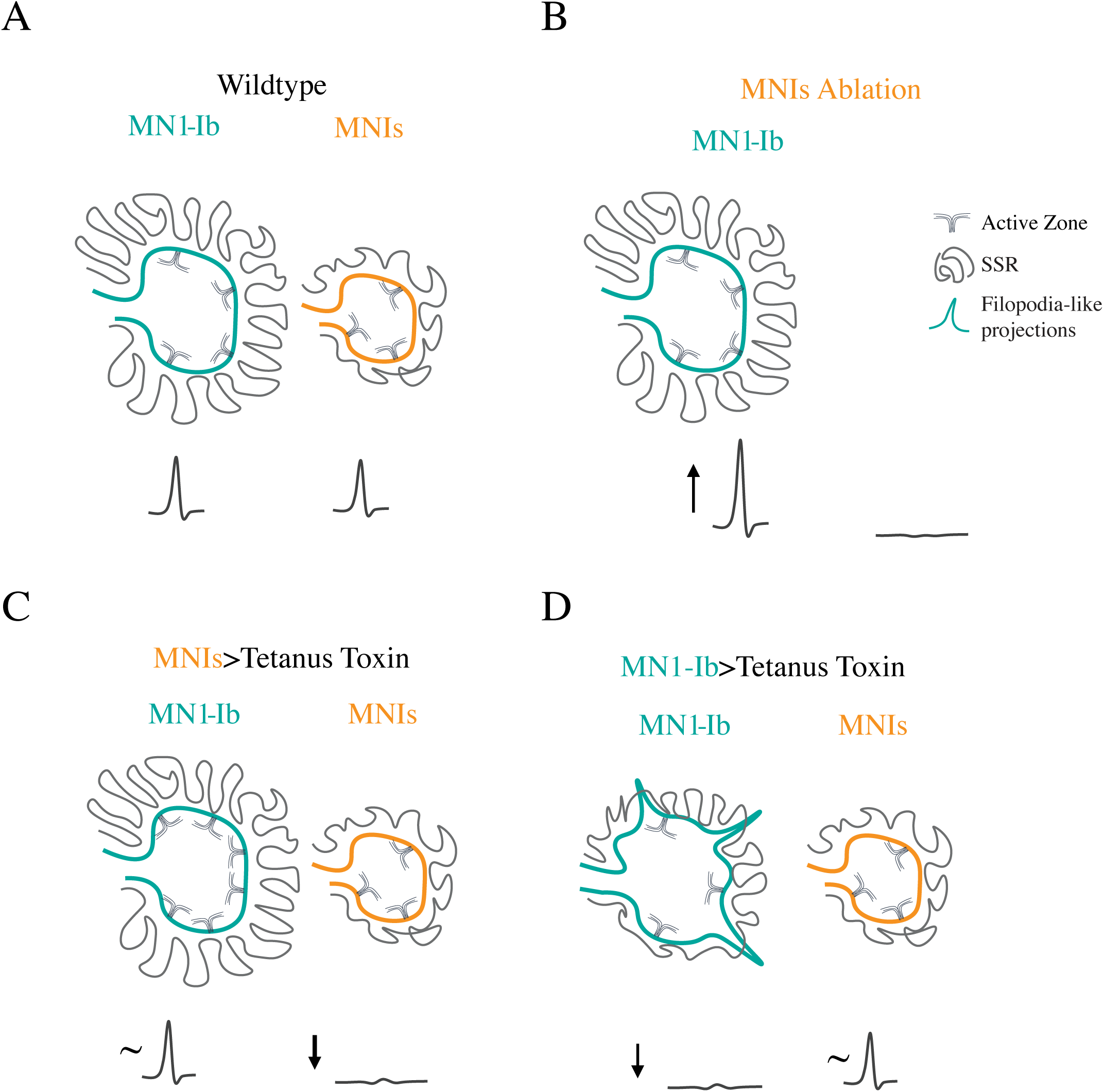
Summary of observed MN1-Ib plasticity. ***A***, In wildtype, MN1-Ib and MNIs provide similar drive to muscle M1. MN1-Ib forms more synaptic boutons and AZs onto M1 compared to MNIs. ***B***, Ablation of MNIs results in increased output from MN1-Ib as evidenced by larger EJPs, but does not trigger increases in bouton or AZ number. ***C***, Silencing of MNIs with tetanus toxin triggers increased bouton and AZ number in the co-innervating MN1-Ib. These changes do not increase presynaptic output from MN1-Ib, with EJP amplitude (∼) at M1 unchanged compared to controls. No structural changes are observed in the silenced MNIs. ***D***, Silencing of MN1-Ib with tetanus toxin results in decreased bouton and AZ number at MN1-Ib terminals. Postsynaptic SSR development is also reduced. Presynaptic filopodia-like projections normally restricted to early 1^st^ instar stage are observed at mature MN1-Ib silenced terminals. No structural or functional (∼) changes occur in the co-innervating MNIs.

The stereotypical connectivity found in the abdominal musculature of Drosophila larvae suggest individual muscles normally allow synaptic innervation from only a single motoneuron of each subclass (Hoang and Chiba, 2001). However, expanded postsynaptic target choice has been observed following muscle loss induced by laser ablation or genetic mutation, with the affected Ib motoneuron targeting inappropriate nearby muscles without altering the innervation pattern of the correctly-targeted Ib neuron (Sink and Whitington, 1991; Keshishian et al., 1994; Chang and Keshishian, 1996). Similarly, ablation of some motoneurons can result in axonal spouting from neighboring connections that target the de-innervated muscle (Chang and Keshishian, 1996). Mis-expression of synaptic cell surface proteins can also alter target choice for some Ib and Is motoneurons (Lin and Goodman, 1994; Kose et al., 1997; Shishido et al., 1998; Ashley et al., 2019). In addition, silencing neuronal activity during development has been demonstrated to induce ectopic NMJs formed primarily by type II neuromodulatory neurons (Keshishian et al., 1994; Jarecki and Keshishian, 1995; White et al., 2001; Lnenicka et al., 2003; Mosca et al., 2005; Carrillo et al., 2010; Vonhoff and Keshishian, 2017). We did not observe any axonal sprouting onto M1 from other motoneurons that resulted in altered target choice when MN1-Ib or MNIs was ablated or silenced in our experiments. Given M1 is the most dorsal muscle of the abdominal musculature, axons from other motoneurons are not present in the direct vicinity, so any signals released from M1 might be insufficient to attract additional innervation. We did find evidence M1 may attempt to promote synaptic innervation when MN1-Ib was silenced with tetanus toxin. Under these conditions, the MN1-Ib terminal maintained an immature-like state with the presence of filopodial-like extensions (Figure 14D). This effect was only observed in Ib motoneurons, highlighting differences in how phasic Is terminals interact with or respond to signals from the muscle. We and others have observed similar filopodial-like extensions at newly forming NMJ connections in late embryos and early 1^st^ instar larvae (Halpern et al., 1991; Broadie and Bate, 1993; Ritzenthaler et al., 2000; Ritzenthaler and Chiba, 2003; Kohsaka and Nose, 2009; Akbergenova et al., 2018). These presynaptic filopodial processes contain elevated levels of the Cacophony (Cac) N-type Ca^2+^ channel and interact with GluRIIA-rich myopodia, with some progressing to form new synapses during early development (Akbergenova et al., 2018). Due to the lack of reinforcement signals caused by absence of synaptic activity in silenced MN1-Ib motoneurons, we hypothesize these processes fail to properly drive AZ assembly and new synapse formation. Indeed, a role for neuronal activity in regulating synaptogenic filopodial stabilization as a precursor to AZ seeding and synapse formation has been recently characterized in the developing Drosophila visual system (Sheng et al., 2018; Özel et al., 2019).

Many forms of plasticity, including synapse elimination at mammalian NMJs, ocular dominance plasticity, and cerebellar climbing fiber pruning, require Hebbian-like input imbalances to trigger synaptic interactions (Wiesel and Hubel, 1963, 1965; Sherman and Spear, 1982; Colman et al., 1997; Sanes and Lichtman, 1999; Walsh and Lichtman, 2003; Turney and Lichtman, 2012; Hashimoto and Kano, 2013; Tomàs et al., 2017; Wilson et al., 2019). As such, we were interested to see if changes in the activity of Ib or Is motoneurons that created an imbalance between the output of the two neurons could drive unique changes compared to when one input was missing. In the case of the tonic Ib neuron this was indeed observed. In the absence of Is input, either due to natural variation in innervation in control animals or following ablation with UAS-RPR, there was no structural response in terms of adding additional release sites. However, loss of Is triggered a functional increase in evoked release from the Ib neuron (Figure 14B). In contrast, when an activity imbalance was created by expressing tetanus toxin in the Is neuron, Ib displayed structural plasticity that increased the number of release sites (Figure 14C). Although the underlying molecular pathways that mediate the two distinct responses are unknown, the results suggest the physical presence of Is likely alters the signaling system(s) responsible for triggering compensation in Ib motoneurons in response to reduced muscle drive.

For every manipulation we made beyond increasing excitability of the neurons, a response from the tonic Ib class was detected, while the phasic Is motoneuron displayed less plasticity. Since each muscle is innervated by only a single Ib motoneuron, plasticity within the tonic subclass may allow more robust and local regulation of muscle function. Although the Is did not show plastic change in response to manipulation of its activity or the co-innervating Ib in our experiments, we cannot rule out that Is neurons are capable of such plasticity but display less sensitivity to putative muscle-derived retrograde signals. In addition, given Is neurons innervate multiple muscles compared to Ib, it is also possible that small plastic changes occurring in Is are not synapse-specific and are distributed over a larger population of AZs onto multiple muscles, resulting in little observed effect at any single postsynaptic target. Similar differences in homeostatic plasticity induction potential in Ib versus Is motoneurons have been previously described following reduced postsynaptic muscle glutamate receptor function, with the Ib motoneuron showing a more robust upregulation of presynaptic release compared to Is, especially in higher extracellular Ca^2+^ (Newman et al., 2017; Li et al., 2018; Cunningham and Littleton, 2019a, 2019b; Genç and Davis, 2019). Together, these results imply tonic Ib motoneurons express distinct plasticity mechanisms that can be triggered by reduced muscle function that are less robust or lacking in the Is phasic subclass.

An important question moving forward is to identify mechanisms that control structural and functional plasticity in tonic Ib motoneurons. Similarly, defining why the Is fails to respond to many of the same manipulations is poorly understood. Whether homeostatic plasticity mechanisms triggered in response to acute or chronic reduction in glutamate receptor function are also activated following the absence or functional silencing of presynaptic inputs as described here is unknown. Several molecular pathways contributing to homeostatic plasticity have been described at the NMJ (Davis, 2006, 2013; Bergquist et al., 2010; Müller et al., 2011, 2012; Müller and Davis, 2012; Younger et al., 2013; Frank, 2014; Wang et al., 2014, 2016; Davis and Müller, 2015; Gaviño et al., 2015; Kiragasi et al., 2017; Li et al., 2018; Ortega et al., 2018; Böhme et al., 2019; Goel et al., 2019; Gratz et al., 2019; Frank et al., 2020). Beyond Drosophila, studies in crustacean motor systems have shown that long-term alterations in activity can induce cell-type specific changes in tonic or phasic motoneuron structure or release properties (Atwood and Wojtowicz, 1986; Lnenicka et al., 1986, 1991; Govind and Walrond, 1989; Lnenicka and Atwood, 1989; Hong and Lnenicka, 1993; Wojtowicz et al., 1994). Given tonic and phasic neurons are abundant in the nervous systems of both invertebrates and vertebrates (Schultz, 2001; Atwood and Karunanithi, 2002; Zucker and Regehr, 2002; Millar and Atwood, 2004; Ventimiglia and Bargmann, 2017), it will be interesting to determine how these two classes interact when innervating a common postsynaptic target in other systems as well. Using GAL4-specific drivers, we are now in a position to characterize the distinct transcriptional profiles of each neuronal subclass to identify candidate mechanisms that mediate the differential plasticity responses of tonic and phasic motoneurons in Drosophila.

## Acknowledgements

This work was supported by NIH grants NS40296 and MH104536 to J.T.L. N.A.S. was supported in part by NIH pre-doctoral training grant T32GM007287. We thank Ellie Heckscher and Gerry Rubin for sharing fly lines. We thank the Bloomington Drosophila Stock Center (Indiana University, Bloomington, IN; NIH P40OD018537), the Drosophila Genome Resource Center (Indiana University, Bloomington, IN; NIH 2P40OD010949-10A1), the TRiP Center at Harvard Medical School (Boston, MA; NIH/NIGMS R01-GM084947), the Developmental Studies Hybridoma Bank (University of Iowa, Iowa City, IA) and the Vienna Drosophila Resource Center (Austria) for providing materials used in this study. We also thank the staff and students in the Cold Spring Harbor Laboratory Drosophila Neurobiology course and members of the Littleton lab for sharing resources and advice.

## Notes

The authors declare no financial or conflicts of interests.

### Competing Interest Statement

The authors have declared no competing interest.

